# Uncovering the embodied dimension of the wandering mind

**DOI:** 10.1101/2024.10.25.620252

**Authors:** Leah Banellis, Niia Nikolova, Malthe Brændholdt, Melina Vejlø, Ignacio Rebollo, Nicolas Legrand, Francesca Fardo, Jonathan Smallwood, Micah G. Allen

## Abstract

When at rest, the mind becomes preoccupied with self-generated thoughts, commonly known as mind-wandering. While the social, autobiographical, and temporal features of these thoughts have been extensively studied, little is known about how frequently the wandering mind turns towards the interoceptive and somatic body. To map this under-explored component of "body-wandering," we conducted a large-scale neuroimaging study in 536 healthy participants, expanding a retrospective multidimensional experience sampling approach to include probes targeting visceral and somatomotor thoughts. Our findings reveal a robust inter-individual dimension of body-wandering characterized by negative affect, high autonomic arousal, and a reduction in socially oriented thoughts. Despite this negative tone, individual differences in the propensity for body-wandering were associated with lower self-reported symptoms of ADHD and depression. Multivariate functional connectivity analyses further revealed that affective, body-oriented thoughts are related to a pattern of thalamocortical connectivity interlinking somatomotor and interoceptive-allostatic cortical networks. Collectively, these results demonstrate that self-generated thoughts exhibit core embodied features which are linked to the ongoing physical and emotional milieu of the visceral body.

**Significance statement:** Neuroimaging studies often treat the resting state as a purely cognitive baseline, overlooking the participant’s embodied experience within the scanner. Here we show that individuals robustly vary in their degree of body-wandering, such that a more somatically focused stream of thought is associated with negative affect, higher arousal, and increased thalamocortical-somatomotor connectivity. Paradoxically, while this state is experienced negatively in the moment, the propensity for embodied mind-wandering correlates with reduced trait level symptoms of depression and ADHD. Collectively, these findings indicate that we must account for the embodied nature of the resting mind to fully understand the neural and clinical mechanisms of self-generated thought.

## Introduction

The human mind ceaselessly shifts between externally driven experiences and self-generated thoughts. The content of this "mind-wandering" is typically characterized along various cognitive dimensions—such as self-reference, temporal orientation, and affect—which map onto unique neural fingerprints and psychopathological traits (Christoff et al., 2016; Mulholland et al., 2023; Smallwood & Schooler, 2015). However, while mind-wandering is traditionally framed as a cognitive process, ongoing thoughts may also be shaped by the body’s fluctuating internal state (Babo-Rebelo et al., 2016; Brown & Herman, 2025; MacVittie et al., 2024, 2025). Visceral rhythms feature centrally in accounts of both emotion and selfhood (Critchley & Garfinkel, 2017; Engelen et al., 2023; Khalsa et al., 2018), yet little is known about how these interoceptive processes integrate within the flow of ongoing thought.

Here, we address this gap by investigating "body-wandering", defined as ongoing thoughts about somato-motor bodily sensations and internal visceral states. To characterize the phenomenological and neurophysiological nature of embodied mind-wandering, we examined individual differences in thought patterns captured in a large cohort during resting-state fMRI. Specifically, we aimed to address three questions: (i) What are the defining mind-wandering content and affective characteristics of body-wandering? (ii) How does this embodied dimension relate to individual differences in mental health and physiological arousal? and (iii) What intrinsic functional brain architecture relates to these body-oriented thoughts?

Ongoing thought varies along fundamental dimensions of content and form, the most common being self-relatedness, temporal focus, and affective tone (Smallwood et al., 2021; Smallwood & Andrews-Hanna, 2013b). These patterns exhibit stable inter-individual differences that map onto distinct neural architectures and clinical profiles (Gorgolewski et al., 2014; Kucyi et al., 2023, 2024). In psychopathology, extreme variations in these dimensions are well-documented: depression is characterized by excessively negative, past-oriented rumination, while ADHD is linked to disruptive attentional lapses and physiological hypo-arousal (Bellato et al., 2023; Hoffmann et al., 2016; Lanier et al., 2021). However, despite the known role of interoception with emotion and selfhood, current models have largely overlooked how the body’s internal state contributes to these intrinsic thought patterns.

Neurobiologically, self-generated thought is largely attributed to the Default Mode Network (DMN) and its dynamic interactions with frontoparietal control and salience networks (Christoff et al., 2016; Hardikar et al., 2024; Turnbull et al., 2019). While specific DMN regions support the social and cognitive features of mind-wandering, the neural basis of body-centric thought remains unclear. Interoception recruits dedicated "allostatic" circuitry spanning ventral salience and somatomotor networks to regulate physiological integrity (Barrett & Simmons, 2015; Kleckner et al., 2017; Zhang et al., 2025). Crucially, fluctuations in arousal modulate default mode network dynamics by recruiting the salience network to gate thalamocortical communication (Groot et al., 2021; Schneider et al., 2016). Insular and cingulate hubs mediate this process by engaging thalamic control pathways that regulate information flow to default mode regions (Allen, 2020; Allen eti al., 2016; Menon & Uddin, 2010). Consequently, we hypothesized that body-wandering emerges from the integration of these systems, characterized by increased connectivity between the canonical networks of ongoing thought and the visceral-sensory architecture responsible for somatic monitoring.

## Results

To investigate the nature of body-wandering and its neural basis, we expanded the well-established Multidimensional Experience Sampling (MDES) framework to include specific probes targeting interoceptive and somato-motor sensations (Martinon et al., 2019; Ruby et al., 2013; Smallwood et al., 2016, 2021; Wang et al., 2018). In total, we used 22 questions addressing social-cognitive, affective, descriptive, and embodied features of ongoing thought (see Table 1). We collected retrospective MDES probe data from 536 participants immediately following a 14-minute resting-state fMRI. Our analysis strategy proceeded in four stages to move from phenomenological description to neural mechanism. First, we established the psychological and physiological properties of body-wandering by examining item-level correlations with affective tone, autonomic arousal, and clinical symptom traits. Second, we applied Exploratory Factor Analysis (EFA) to identify the latent dimensions that organize these varying thought patterns. Third, we utilized Canonical Correlation Analysis (CCA) to uncover the multivariate relationships between these thought patterns and whole-brain functional connectivity signatures. Finally, we validated the convergence between the resulting neural signatures and the latent psychological factors and situated the findings within the principal gradients of functional brain organization.

**Table 1:** Multidimensional interoceptive experience sampling items asked after resting state fMRI.

| Dimension | Item | Scale=0 | Scale=100 |
| --- | --- | --- | --- |
| Future | My thoughts involved future events. | Not at all | Completely |
| Past | My thoughts involved past events. | Not at all | Completely |
| Self | My thoughts involved myself. | Not at all | Completely |
| Other | My thoughts involved other people. | Not at all | Completely |
| Positive | My thoughts were positive. | Not at all | Completely |
| Negative | My thoughts were negative. | Not at all | Completely |
| Words | My thoughts were in the form of words. | Not at all | Completely |
| Vivid | My thoughts were vivid as if I was there. | Not at all | Completely |
| Vague | My thoughts were detailed and specific. | Not at all | Completely |
| Spontaneous | My thoughts were: | Spontaneous | Deliberate |
| Focus | My thoughts were focused. | Not at all | Completely |
| Repetitive | My thoughts were repetitive. | Not at all | Completely |
| Temporal distance | My thoughts were related to: | The here and now | A more distant time |
| Images | My thoughts were in the form of images. | Not at all | Completely |
| Arousal | In general, I felt more: | Sleepy | Alert |
| Body | My thoughts were about my body. | Not at all | Completely |
| Breath | My thoughts were about my breathing. | Not at all | Completely |
| Heart | My thoughts were about my heartbeat. | Not at all | Completely |
| Movement | My thoughts were about moving. | Not at all | Completely |
| Bladder | My thoughts were about my bladder. | Not at all | Completely |
| Skin | My thoughts were about my skin. | Not at all | Completely |
| Stomach | My thoughts were about my stomach. | Not at all | Completely |

### Identifying relationships of bodily-related content to ongoing thought patterns

We compared the overall prevalence of body-focused versus cognitive content by aggregating the ratings of the 8 newly added interoceptive and somatomotor items and contrasting them with the average of the 14 original MDES items. This analysis revealed that interoceptive thoughts were reported significantly less frequent than typical cognitive, affective, or descriptive themes (Body Average Median = 36.3 vs. Cognitive Themes Average Median = 47; *W*(535) = 113236.5, p < 0.001, Rank-Biserial correlation (rB) = 0.574). Within the somatic domain, broader thoughts about movement and the body in general were significantly more prevalent than sensations of specific organs, while breathing emerged as the dominant interoceptive feature (Figure 1A, see Supplementary Table 1 for full statistics).

**Figure 1:**
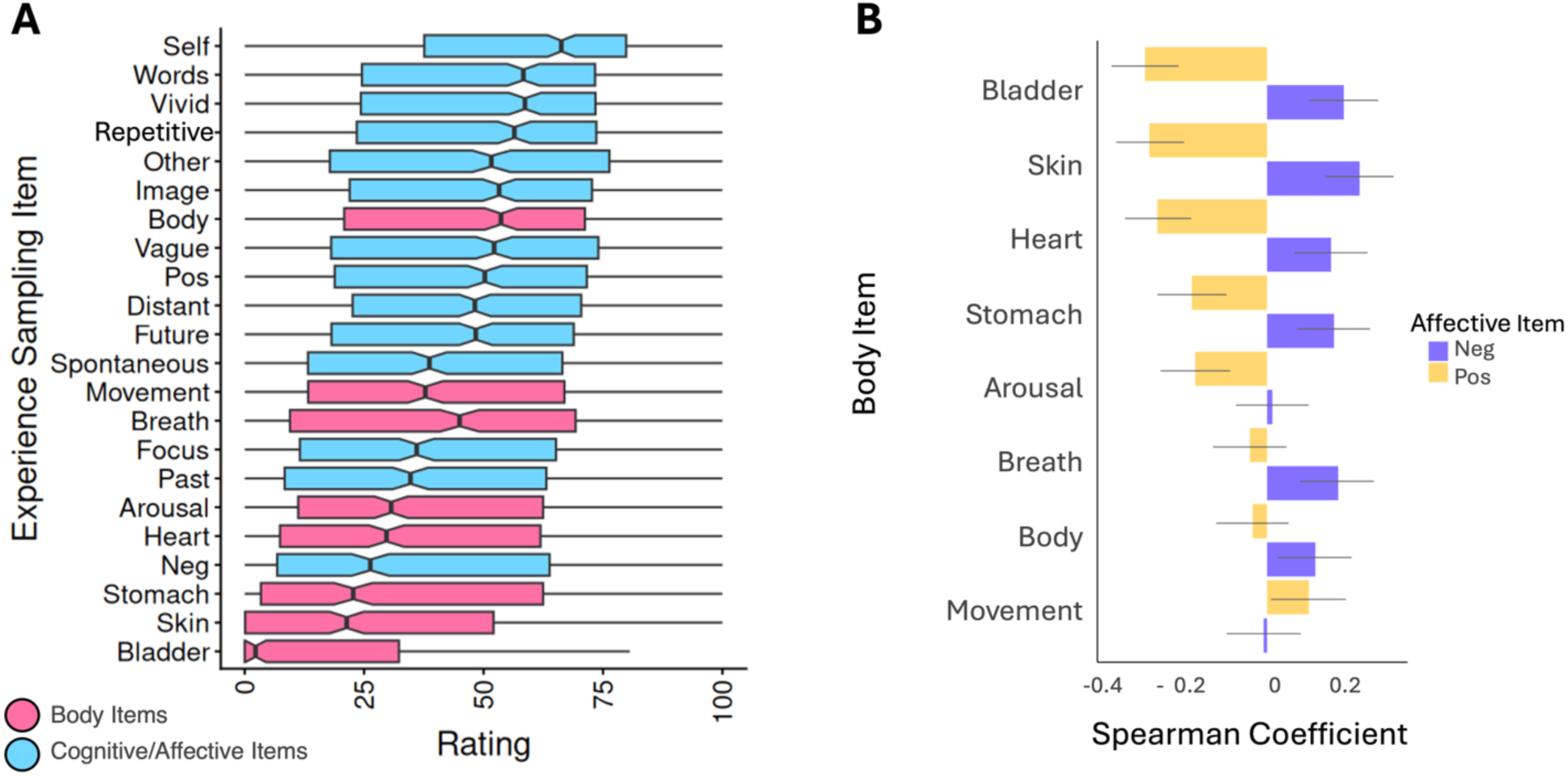
Ongoing thought item ratings including body-related, cognitive, and affective content after resting state fMRI. A) Boxplots of mind-wandering item ratings (pink: body-related items and blue: classical cognitive/affective/descriptive MDES items). B) Spearman correlation analysis of affect and body-related mind-wandering items (yellow: body-related items correlations with the positive mind-wandering item, purple: body-related items correlations with the negative mind-wandering item, error bars reflect bootstrap 95% confidence intervals via 10’000 resamples. See Supplementary Table 2 for full FDR-corrected correlation statistics).

While resting thoughts were generally positive in valence (positive-negative difference = 15.820, W(535) = 86585, p < 0.001), we observed substantial individual variability in affective tone (Range = −100 to 100). Examining the constituent components of this individual variation revealed that interoceptive body-wandering is substantially linked to heightened negative emotion. The majority of items probing visceral and somatic sensations, and in particular the heart, bladder, skin, and stomach, were consistently correlated with both reduced positive and heightened negative content (all pFDR =< 0.001, see Figure 1B and Supplementary Table 2). Thus, unlike the typically positive drift of mind-wandering, attention to the body during resting state fMRI is characterized by a more negative affective thought pattern.

Compatible with this, we found that traditional cognitive and descriptive features of ongoing thought, such as temporal focus, thought focus, and self vs social reference, were generally associated with increased positive emotional tone (all pFDR < 0.001). These items correlated with greater positive and/or reduced negative affect (see Supplementary Figure 1). Together, these results establish a fundamental affective divergence: whereas a more cognitive mind-wandering style tends to be associated with more positive thoughts, a mind-wandering style defined by interoceptive thoughts is characteristically more negative.

To verify the stability of these phenomenological patterns, we performed a series of robustness checks. First, we confirmed that all key associations persisted when controlling for potential confounds, including age, gender, and body mass index (BMI) (see Supplementary Figure 1). Second, we performed a split-half analysis between odd and even participants (N = 268 each), which revealed excellent intraclass correlations for item prevalence ratings (ICC (A,1) = 0.937, p < .001), and cognitive-affect relationships (ICC = 0.907, p < 0.001), as well as good robustness for the intraclass correlation profiles of the body-affect associations (ICC = 0.882, p < .001). This confirms that the prevalence, organization, and affective divergence of body-wandering are robust, reproducible features of the resting mind.

### Body-related mind-wandering is linked to reduced ADHD and depression symptoms

We assessed individual mental health symptoms using the Adult ADHD Self-Report Scale (ASRS) (Kessler et al., 2005) and the Major Depression Inventory (MDI) (Bech et al., 2015). Correlating these scores with the expanded MDES item set revealed that the tendency to think about the body was associated with reduced psychopathology (see Figure 2A and 2C). Specifically, higher ratings of thoughts related to the heart, bladder, and skin were linked to significantly lower symptom severity for both ADHD (*r*_s_ range = −0.104 to −0.211, pFDR < 0.05) and depression (*r*_s_ range = - 0.105 to −0.149, pFDR < 0.05). Conversely, depression was characterized by greater engagement in mental time travel, exhibiting positive correlations with the past and future (*r*_s_ range = 0.123 to 0.164, pFDR < 0.05). These associations remained robust after controlling for age, gender, and BMI (Supplementary Figure 2; see Supplementary Table 3 for full cross-correlation statistics), were similar when estimated using latent variable analyses (see Supplementary Figure 3), and showed good reliability as demonstrated by split-half intraclass correlation (ICC_(A,1)_ = 0.861, p < 0.001). Collectively, these results suggest that a reduction in spontaneous interoceptive thought is a shared correlative marker of both mood and attentional dysfunctions, while a specific detachment from the present moment via temporal rumination is characteristic of depressive symptomatology. For full inventory scoring details, see Methods; see Supplementary Table 4 for individual survey items.

**Figure 2:**
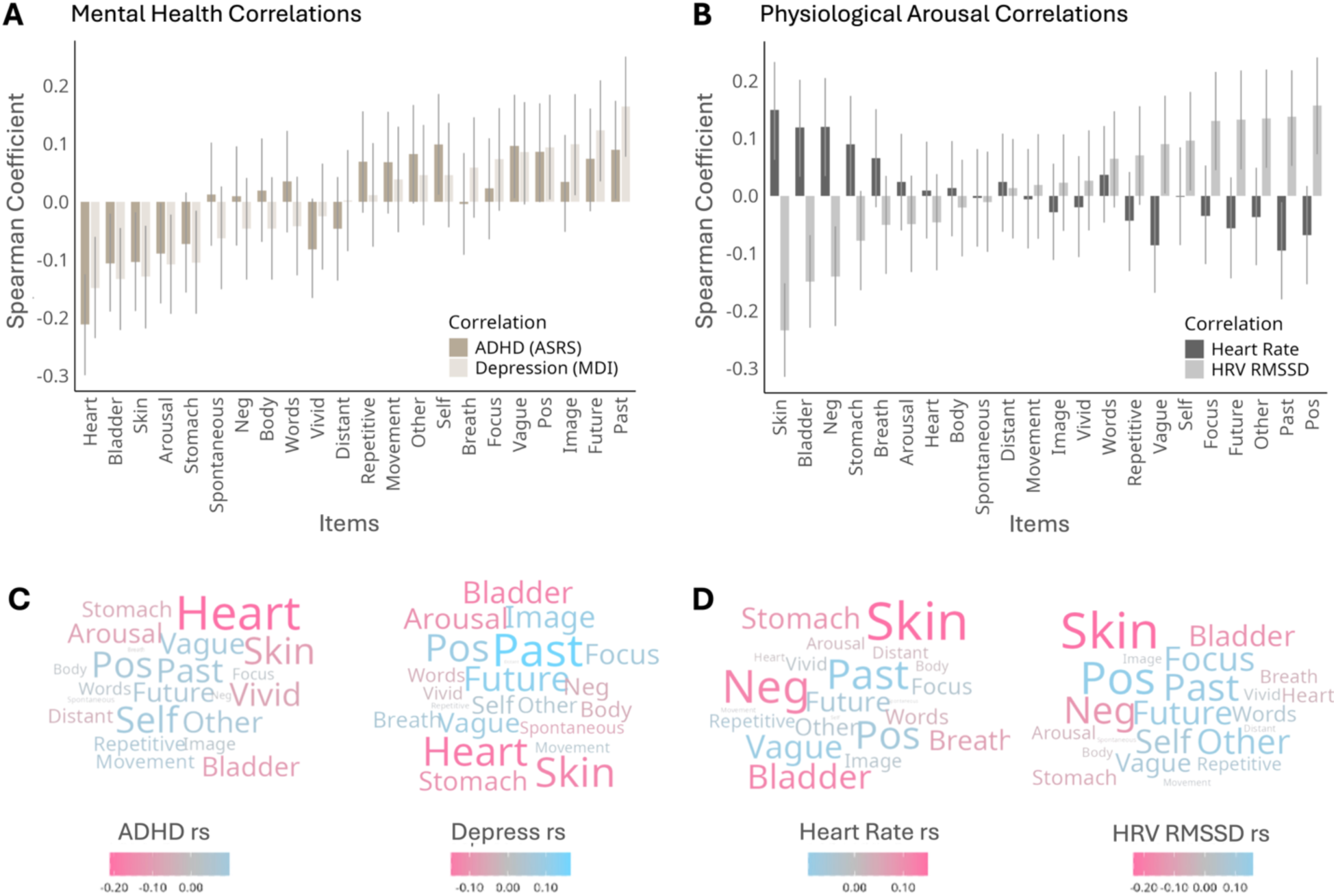
Body-related and cognitive thought patterns exhibit opposing associations with mental health and physiology. (A and C) Spearman correlations revealing that body-wandering items align with reduced symptom severity for ADHD and depression (ASRS and MDI sum scores respectively), whereas social-cognitive and temporal items are associated with higher symptom severity. (B and D) Physiologically, increased somatic and interoceptive thoughts are characterized by markers of high autonomic arousal (positive correlations with Heart Rate; negative with HRV-RMSSD), while cognitive aspects of mind-wandering reflect a low-arousal state (positive parasympathetic HRV-RMSSD correlations) (see Supplementary Table 3 for full FDR-corrected statistics). These item-level patterns mirror the relationships observed in latent variable analyses (see Supplementary Figure 3). Error bars reflect bootstrap 95% confidence intervals via 10’000 resamples.

### Autonomic arousal correlates with interoceptive thought patterns

We found that interoceptive and somatic thoughts, in particular regarding the skin and bladder were correlated with individual cardiac markers of high autonomic arousal, including elevated heart rate (*r*_s_ range = 0.12–0.15; all pFDR < 0.02) and reduced parasympathetic markers of heart rate variability (RMSSD; *r*_s_ range = −0.15 to −0.23; all pFDR < 0.005) (see Figure 2B and 2D). Conversely, more cognitive contents (e.g., future, past, other, focus) were associated with increased parasympathetic heart rate variability (*r*_s_ range = 0.13–0.14; all pFDR < 0.01), reflecting lower physiological arousal. Affective tone showed a similar physiological divergence: negative affect was associated with elevated heart rate (*r*_s_ = 0.12) and reduced variability (*r*_s_ = −0.14), while positive affect was linked to increased heart rate variability (*r*_s_ = 0.16; all pFDR < 0.02) (see Supplementary Table 3 for full statistics). Complementary patterns were observed in respiratory and electrogastrographic metrics, where somatic thoughts similarly tracked markers of heightened arousal while cognitive wandering tracked with reduced arousal (see *S*upplementary Figure 4 and 5). These associations remained consistent when controlling for age, gender, and BMI (see *S*upplementary Figure 2) and demonstrated good reliability across data splits (*ICC*_(A,1)_ = 0.834, *p* < 0.001) and were also similar when using latent variable analyses (see Supplementary Figure 3).

### Uncovering the latent embodied structure of self-generated thought

To determine how bodily themes organize within the broader landscape of mental experience, we applied Exploratory Factor Analysis (EFA) to the expanded MDES item set. This analysis uncovered a robust affective, embodied dimension of thought defined by the contrast between physiological and social content, and the overall valence of thoughts (Factor 1; 9% variance). This factor loaded heavily on thoughts related to interoceptive and somatic sensations, specifically skin, heart, bladder, and stomach, alongside high negative affect. In the opposite direction, this dimension showed strong negative loadings for positive affect and social-cognitive thoughts (self, other). This polarity suggests that thoughts about the body do not occur in isolation but rather form part of a specific somato-affective trait that competes at an individual level with pleasant, socially oriented thoughts. In contrast, a second independent factor (Factor 2; 8% variance) captured the cognitive contents of mind-wandering, clustering mental time travel (past, future) with features of clarity and form (focus, image, vague, repetitive) (see Figure 3 and Supplementary Figure 6 for full pattern coefficients). These dimensions demonstrated high and moderate structural stability across data splits, respectively (Factor 1: *r* = 0.92; Factor 2: *r* = 0.68)

**Figure 3:**
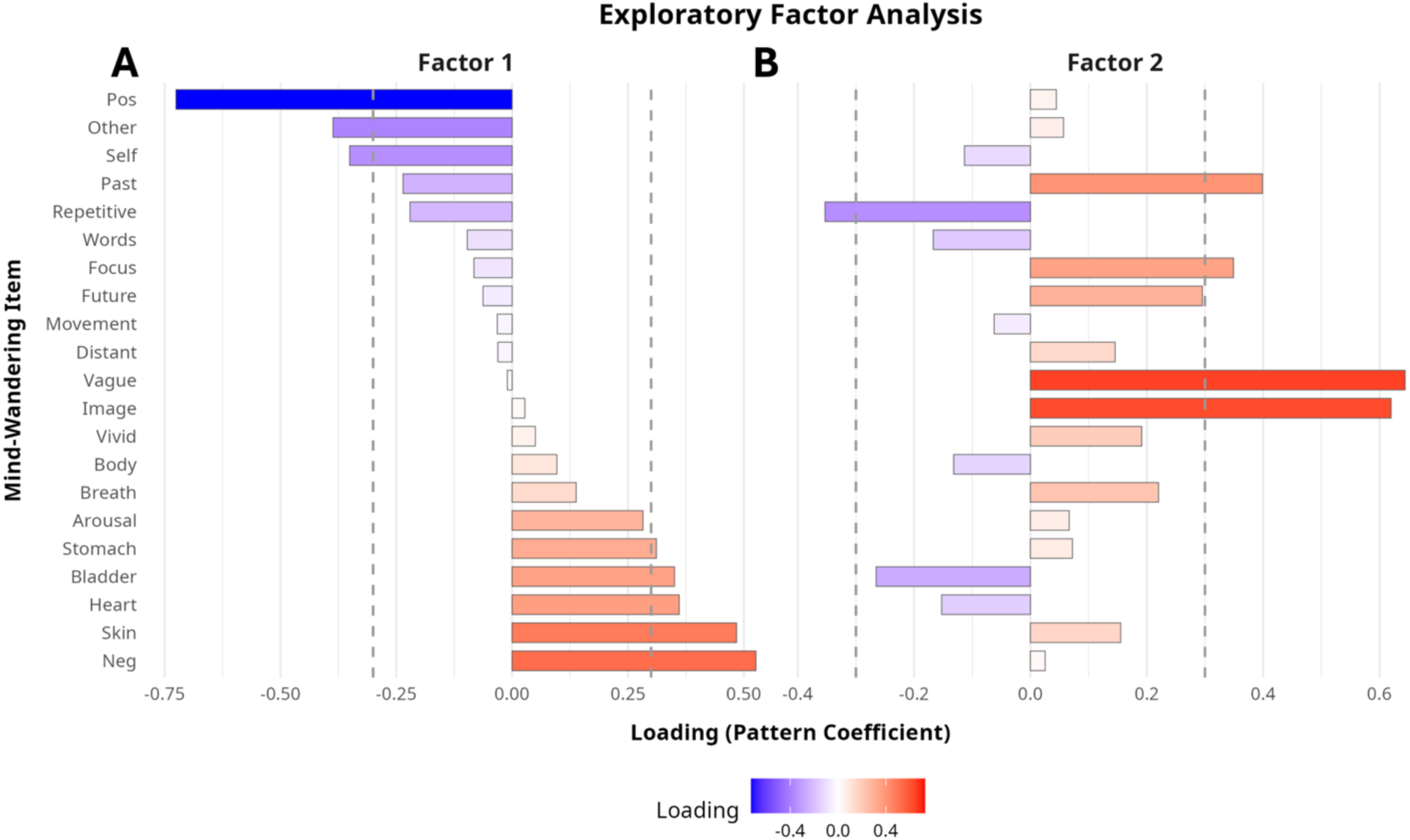
The organization of self-generated thought reveals a trade-off between affective valence, interoceptive, and social-cognitive content. Exploratory Factor Analysis (EFA) identifies two principal dimensions of the resting mind. Factor 1 captures a somato-affective dimension defined by the polarization between visceral, negative-valenced thoughts (positive loadings; red) and pleasant, socially-oriented mentation (negative loadings; blue). Factor 2 reflects the cognitive modality of thought experience, organizing mental time travel (*Past*, *Future*) alongside features of form (*Image*, *Focus, Vivid, Repetitive*). Dashed lines indicate a loading threshold of > absolute 0.3. See Supplementary Figure 6 for full pattern coefficients.

### Intrinsic functional connectivity signature of visceral-affective thought

We further investigated the intrinsic functional architecture that supports body-related thoughts. To do so, we employed cross-validated Canonical Correlation Analysis (CCA) to identify latent dimensions that maximize the correspondence between individual differences in reported thoughts and functional brain connectivity profiles as measured across 216 cortical and subcortical brain parcels (see Methods). This analysis revealed a single, significant, and cross-validated mode of neuro-cognitive covariation (canonical variate out-of-sample *r* = 0.35, *p* = 0.001) that links affective bodily thoughts to a subcortical-somatomotor brain connectivity profile (see Figure 4 and Supplementary Table 5).

**Figure 4:**
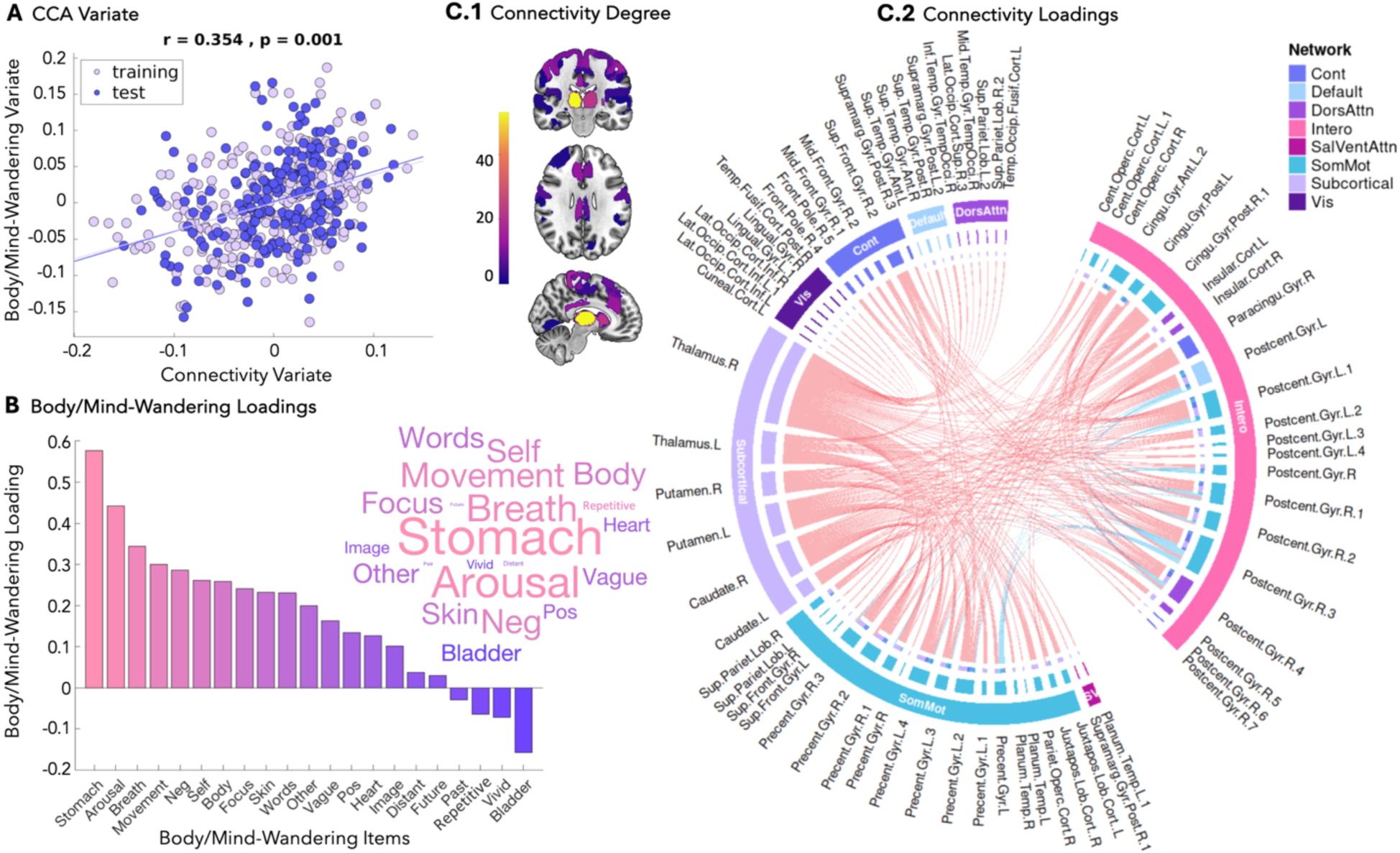
The connectomic fingerprint of embodied-affective mind-wandering. (A) Canonical Correlation Analysis (CCA) revealing a significant, cross-validated mode of covariation between ongoing thought and functional connectivity (out-of-sample *r* = 0.35, *p* = 0.001). Each dot represents a participant. (B) The phenomenological loadings of individual thought patterns on the canonical mode. Positive loadings define a dimension anchored by visceral sensations (stomach, arousal, breath), movement, negative affect, and self-orientation. (C) The intrinsic connectivity architecture supporting this dimension, thresholded at the top 1% of canonical loadings (edge weights > absolute 0.55). (C.1) Nodal degree of the high-loading edges, identifying the right thalamus as the network’s primary hub (*N*=53 edges), followed by the right postcentral gyrus (24 edges) and striatum (caudate/putamen). (C.2) Chord plot illustrating the strongest functional connections. The network is dominated by integration between Subcortical (purple) and Somatomotor (blue) systems. *Note:* For visualization, regions implicated in interoceptive allostasis (insula, cingulate, postcentral gyrus, central opercular cortex, and paracingulate cortices) were grouped into a "Interoceptive" cluster (pink) (Amassian, 1951; Azzalini et al., 2019; Downman, 1951; Engelen et al., 2023); inner ring colors denote their original Yeo-7 network assignments. The CCA demonstrated a significant relationship between the training and test set in 4/5 of the data folds (80%), revealing high robustness and generalisability across the data folds.

Phenomenologically, this analysis identified a dimension of self-generated thought predominately defined by embodied thought. The multivariate pattern was most strongly characterized by interoceptive and somatic items: specifically, thoughts about the stomach (loading = 0.58), subjective arousal (0.44), breathing (0.35), and movement (0.30) made the greatest contributions to the canonical mode. We further observed moderate loadings for negative affect (0.29) and self-focus (0.26) (see Figure 4B). Notably, unlike the bipolar "Body vs. Social" structure observed in the exploratory factor analysis, this brain-constrained mode did not exhibit a strong opposing pole; thoughts about others (0.20) and positive affect (0.14) showed weaker, positive associations. Consequently, this complementary neural signature maps the tendency to engage in high-arousal embodied thought, prioritizing a specific mode of visceral thoughts, negative affect, and self-reflective details.

In the resting connectome, this dimension is related to a highly integrated network centered on somatomotor, allostatic, and subcortical systems. The neural signature was predominately characterized by heightened functional coupling between the thalamus and striatum (caudate/putamen) and the primary somatosensory and motor cortices (see Figure 4C). Specifically, the connectivity edges with the highest canonical loadings linked the right thalamus and right caudate directly to the bilateral precentral and postcentral gyri (top 1% loadings > 0.55; see Supplementary Table 6). Beyond these sensorimotor hubs, the mode also recruited regions critical for interoceptive allostasis, including the insula, cingulate, and paracingulate gyri. The right thalamus emerged as the most highly connected hub within this array (degree = 53 edges; see Figure 4C.1), suggesting a prominent topological role within the network linking visceral and somatic regions during the stream of ongoing thought (see Supplementary Table 7). Positive loadings were most prevalent (red chord-plot connections of Figure 4C.2), indicating that increased subcortical-somatomotor connectivity is primarily involved in this embodied thought integration.

### Macroscale Organization of Body-Wandering

To situate the body-wandering signature within the broader landscape of cortical organization, we assessed the correspondence between the principal CCA mode and established functional gradients of the human brain (Margulies et al., 2016). We found that the topography of body-wandering is significantly associated with Gradient 1, the principal axis differentiating unimodal sensory-motor regions from transmodal association cortex (*p* < 0.001, spin permutation test with 2500 permutations (Alexander-Bloch et al., 2018)). Specifically, the negative poles of the body-wandering mode align with the gradient’s somatomotor anchor in the postcentral and precentral gyri, while its positive poles track the transmodal apex in the right superior/middle frontal and bilateral cingulate cortices (Fig. 5). This indicates that body-focused thought bridges the hierarchy between immediate somatic sensation and abstract self-referential processing. This alignment was specific to the principal gradient; associations with the visual-to-motor axis (Gradient 2) did not survive correction for multiple comparisons (*p* = 0.048; Bonferroni α = 0.01), and no significant relationships were observed for Gradients 3–5 (smallest *p* = 0.189).

**Figure 5:**
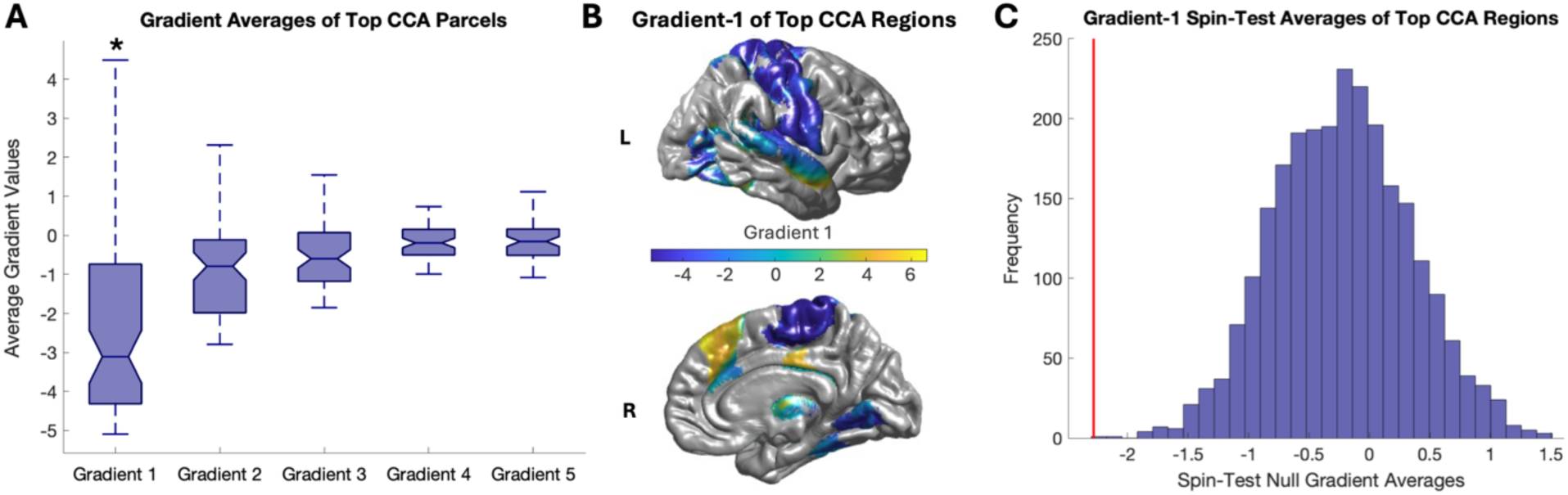
Relationship of the top 1% of brain regions from the affective body-wandering CCA with gradient brain variation dimensions via spin permutation tests. A) Boxplots of average values from each gradient atlas (Margulies et al., 2016) for each of the top 1% CCA brain parcels (asterisks mark whether significant via a spin-test null distribution with 2500 permutations). (B) The top 1% of CCA brain regions coloured by the Gradient-1 atlas, including high negative loadings for somatosensory and motor regions (postcentral and precentral gyrus), and high positive loadings for right frontal (i.e., right superior and middle frontal gyrus) and bilateral cingulate regions. (C) Spin-test result for the average Gradient-1 atlas value across the top 1% of brain parcels (red vertical line), in comparison to the average values determined from a spin-test null distribution with 2500 permutations (Alexander-Bloch et al., 2018).

### Convergent Validity of the Body-Wandering Connectome

Finally, we sought to establish the convergent validity of the derived embodied dimension of thought across our analytical approaches. Across item-level correlations, EFA, and CCA, we consistently observed a clustering of somatic thoughts with negative affect. However, the precise composition of this embodied-affective state varied by method: while the unsupervised EFA emphasized cardiac, stomach, skin and bladder sensations as opposing positive social thought, the brain-constrained CCA mode prioritized gastric, respiratory, and arousal sensations alongside a moderate involvement of self-related thought.

To determine if these variations reflect distinct phenomena or at least partially overlapping constructs, we tested whether the brain network supporting the unsupervised behavioral factor (Factor 1) overlapped with the network identified by the CCA. We conducted a mass-univariate regression across all 23,220 functional connectivity edges to identify brain regions associated with the individual EFA factor scores, with significance determined via non-parametric permutation testing (see Figure 6).

**Figure 6:**
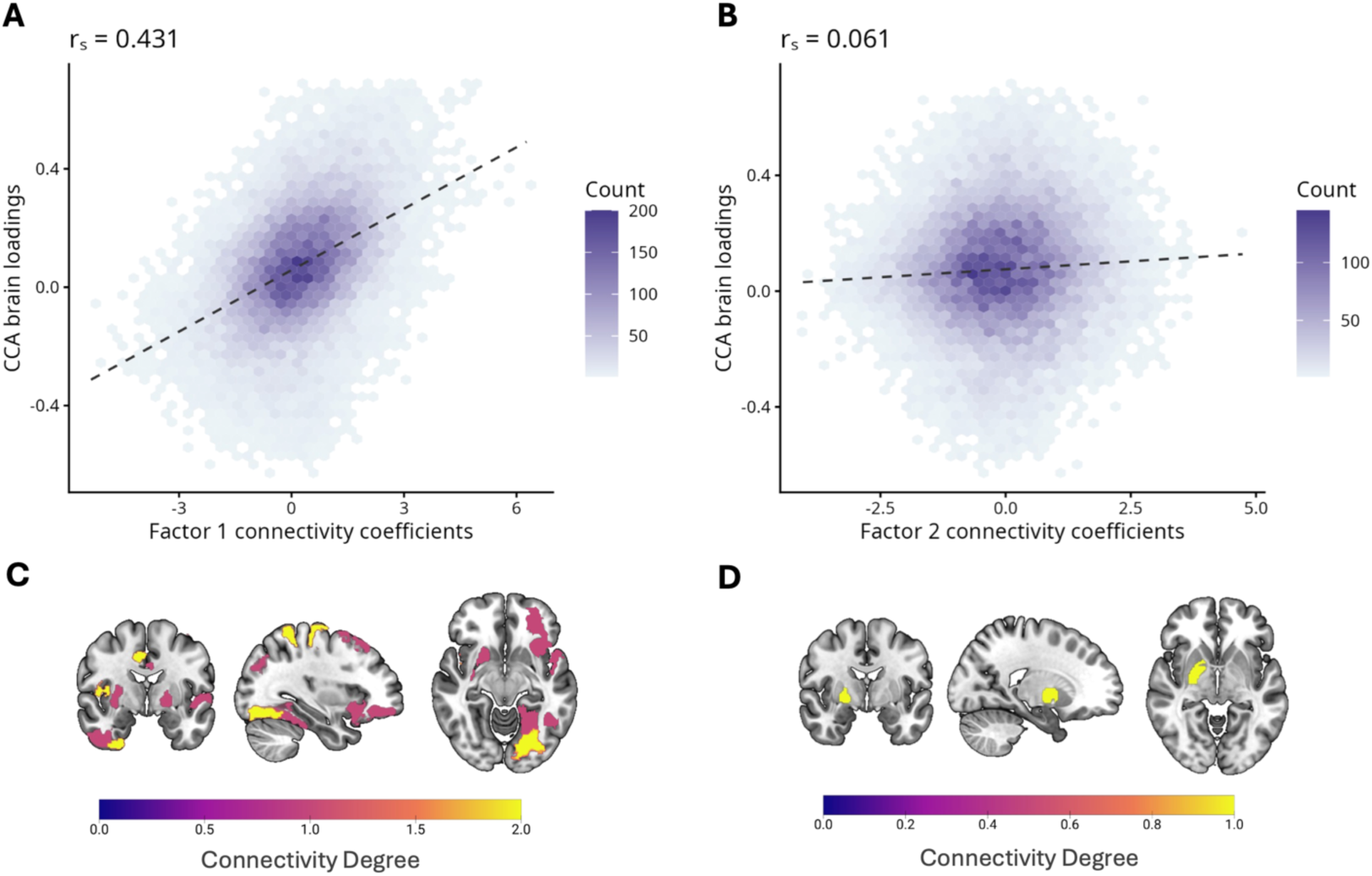
Convergent validity of the body-wandering connectome. (A and B) Hexagonal binned plots displaying the spatial density of correlations between the multivariate CCA connectivity profile (x-axis) and mass-univariate EFA connectivity regression coefficients (y-axis) for (A) the Body-Wandering factor (*r* = 0.431, p < 0.001) and (B) the Mental Time Travel factor (*r* = 0.061). The strong correlation in (A) supports that the supervised CCA and unsupervised EFA converge on the same brain connectivity substrate. (C and D) Node degree distributions (sum of significant edges for each brain region) of edges significantly associated with (C) Body-Wandering (17 edges; predominantly somatomotor, salience, and visual networks) and (D) Mental Time Travel (1 edge) (see Supplementary Table 8). Significance was assessed via non-parametric permutation testing (10’000 permutations). Note, given the large number of connectivity edges (>23,000), even very small correlations are expected to reach statistical significance; interpretation therefore focuses on effect size and network specificity.

This analysis revealed a robust convergence across the estimated dimensions. The spatial profile of functional connectivity predicting the EFA-derived body-wandering factor was significantly correlated with the connectivity weights of the CCA mode (r = 0.431, p < 0.001; see Figure 6A and 6C). Specifically, the EFA factor exhibited a comparable topographical organization, recruiting a similar constellation of edges across somatomotor, salience, and visual networks (see Supplementary Table 8). In contrast, the "mental time travel" descriptive dimension (Factor 2) showed negligible neural overlap with this architecture (*r* = 0.061, p < 0.001). This supports that despite the nuanced re-weighting of specific interoceptive features, when constraining for brain connectivity, both analyses tap into an overlapping neural dimension.

### Control Analyses

We performed additional control analyses to ensure our findings were not driven by potential confounds such as state-based bladder-related physical discomfort, trait-level somatic distress, or trait anxiety individual differences. To this end, we controlled for these factors in both the psychophysiological correlation analysis and the brain connectivity CCA by regressing out specific nuisance variables from the mind-wandering responses prior to analysis. This procedure was performed separately for momentary bladder sensation (using individual ratings for the ‘Bladder’ item), trait-level somatic distress (using the Patient Health Questionnaire somatic symptoms module, PHQ-15 (Kroenke et al., 2002)), and trait anxiety (using the State-Trait Anxiety Inventory, STAI-Trait (Spielberger, 1983)). We observed results highly similar to our main findings across all controls; both the psychophysiological associations (see Supplementary Figures 7-12) and the brain connectivity neural signatures (see Supplementary Figures 13-15) remained significant and robust after correction (although there are some minor variation in the significance of individual item psychophysiological correlations; see Supplementary Figures 7-12). These results suggest that the identified body-wandering phenotype is not driven by individual differences in bladder discomfort, general somatic distress, or trait anxiety.

## Discussion

Research on mind-wandering has primarily focused on the balance between internally and externally directed attention, with emphasis on autobiographical, temporal, and social axes of thought. This focus has overlooked a key implication of internally oriented thought: the body, as a primary anchor of selfhood, provides continuous information about the organism’s physiological state and its ongoing regulation. We here identified *body-wandering* as an embodied dimension of self-generated thought, extending prevailing accounts of mind-wandering to include visceral and somatomotor experience. Our findings identify a robust inter-individual tendency toward body-oriented mentation, linked to autonomic arousal, affective tone, and intrinsic thalamo-somatomotor connectivity.

### Body-wandering as an inter-individual feature of thought

We assessed spontaneous thought using retrospective experience sampling following a single 14-minute resting-state scan. Accordingly, the present results index stable inter-individual differences in the relative salience of bodily versus cognitive thought contents during rest, rather than the moment-to-moment dynamics of thought. This distinction is important, given evidence that spontaneous thoughts fluctuate on short time scales and that interoceptive attention and internal cognition can compete dynamically at the state level (Brown & Herman, 2025; Groot et al., 2021; Kucyi et al., 2023; Vanhaudenhuyse et al., 2011). The findings therefore speak to dispositional tendencies in how individuals experience rest, consistent with prior work demonstrating that individual differences in ongoing thought are reliable and systematically related to physiology and intrinsic brain organization (Kucyi et al., 2024; Wang et al., 2018). It is also important to note that being inside an MRI scanner is itself an unusual physical experience. Lying motionless in a narrow tube with limited sensory input likely shapes both bodily state and ongoing experience and thus provides an important context for interpreting resting-state brain imaging studies.

### Embodied relationship with affective tone and autonomic arousal

The present findings demonstrate that the bodily contents of self-generated thought constitute a particularly affectively charged dimension, extending prior work on emotion in mind-wandering. Across item-level analyses, latent structure, and multivariate brain–behavior coupling, body-wandering was consistently associated with more negative affect. Thoughts about visceral and somatic sensations—particularly related to the heart, stomach, skin, and bladder —were linked to reduced positive and increased negative affect, whereas socially oriented and descriptive cognitive features showed the opposite pattern. This profile aligns with recent work linking interoceptive attention to negative (Brown & Herman, 2025; Poerio et al., 2024), while contrasting with affective patterns observed to co-occur with social and descriptive thought content (Ruby et al., 2013; Wang et al., 2018). This dissociation extends content-based accounts of mind-wandering, which emphasize that the emotional consequences of spontaneous thought depend on its content and context rather than on its mere occurrence (Andrews-Hanna et al., 2014; Mason et al., 2013; Smallwood & Andrews-Hanna, 2013b; Smallwood & Schooler, 2015).

Body-wandering was also linked to variation in autonomic physiology. Individuals reporting more bodily focused thoughts exhibited higher heart rate and reduced parasympathetic heart rate variability, with convergent patterns across respiratory and gastric measures. In contrast, cognitively oriented mind-wandering tracked lower-arousal physiological states. These results refine earlier observations linking dysphoric or perseverative mind-wandering to heightened autonomic activation (Ottaviani et al., 2015; Smallwood et al., 2007), showing that elevated arousal may be tied to embodied thought content rather than to spontaneous thought more generally.

This pattern is also compatible with work implicating the locus coeruleus–norepinephrine system, a central regulator of arousal, in mind-wandering. Prior studies have linked elevated tonic arousal to internally oriented processing and perceptual decoupling (Groot et al., 2021; Mittner et al., 2016; Smallwood et al., 2021). The present observation of differential associations between arousal and embodied versus cognitive thought content complements these accounts and suggests that heightened arousal may bias mind-wandering toward interoceptively salient signals, potentially via noradrenergic modulation of neural gain across these channels.

### Neural signature and convergent validity

At the neural level, affective body-wandering was associated with a characteristic pattern of intrinsic connectivity linking thalamic and striatal regions with primary somatosensory and motor cortices, alongside insular and cingulate areas implicated in interoceptive and allostatic processing (Engelen et al., 2023; Kleckner et al., 2017; Menon & Uddin, 2010; Zhang et al., 2025). Critically, the connectivity profile associated with the exploratory somato-affective factor showed strong spatial correspondence with the brain-constrained multivariate solution, whereas the factor capturing temporal and descriptive features of thought showed minimal overlap. This convergence demonstrates that the identified thalamocortical–somatomotor architecture reflects a consistent connectivity profile of embodied spontaneous thought rather than an artifact of a particular analytic choice.

Recent work has shown that spontaneous thought elicited via a free association word task engages bodily representations in a content-specific manner, with body maps predicting trial-wise affective valence and self-relevance and recruiting default-mode, limbic, and somatosensory regions (Shin et al., 2023). Whereas these findings emphasize the representational mapping of bodily signals onto cortical regions during within-subject individual thoughts, the present CCA identifies a distributed somatomotor–subcortical connectivity architecture associated with an inter-individual tendency toward embodied spontaneous thought.

This organization differs from canonical descriptions of mind-wandering emphasizing default-mode and frontoparietal control networks (Andrews-Hanna et al., 2014; Smallwood et al., 2012). Rather than reflecting sensorimotor disengagement (Groot et al., 2021; Smallwood et al., 2011; Wang et al., 2018), body-wandering appears to involve sensorimotor and subcortical circuitry during internal experience. Consistent with this interpretation, the spatial distribution of this network aligns with the principal unimodal–transmodal gradient, bridging somatomotor regions and higher-order self-referential frontal and cingulate cortices (Margulies et al., 2016).

### Clinical associations and implications

Despite its negative affective tone and elevated arousal, individual differences in body-wandering were associated with lower self-reported symptoms of depression and ADHD. In contrast, depressive symptoms were characterized by greater temporal displacement toward past and future thought, consistent with extensive evidence linking depression to abstract, temporally decoupled rumination (Andrews-Hanna et al., 2014; Killingsworth & Gilbert, 2010; Kucyi et al., 2023; Smallwood et al., 2007). These findings argue against a simple equivalence between negative affect during rest and maladaptive cognition.

One speculative interpretation is that spontaneous attention to bodily sensations, while often experienced negatively in the moment, reflects preserved engagement with ongoing sensory input. Such embodied engagement may be protective relative to forms of mind-wandering that are detached from the current bodily state. This view aligns with recent work showing dissociable trait/state relationships between interoception and internal thought: while trait-level interoceptive attention is linked to anxiety-related patterns of negative thinking, state-level attention to the body competes with internal thought (Brown & Herman, 2025). Furthermore, ecological evidence links body awareness to the maintenance of a first-person perspective in daily life (MacVittie et al., 2024). Importantly, these associations persisted after controlling for trait anxiety and somatic symptom burden.

### Limitations and future directions

Several limitations should be acknowledged. First, the retrospective nature of experience sampling precludes inference about causal directionality between bodily state, arousal, and thought content. Future research should explore the dynamical interaction between thoughts about the body and cognitive features over time (Coulborn & Fernández-Espejo, 2022; Martinon et al., 2019; Vanhaudenhuyse et al., 2011). Second, although extensive control analyses argue against trait anxiety or somatic symptoms as primary drivers, momentary interoceptive discomfort in the scanner such as from bladder-related sensations may contribute to variance in a subset of individuals. When controlling for bladder item ratings, this was apparent in some weaker associations, while the overall psychophysiological patterns and brain–body connectivity signatures remain robust. Future studies employing more detailed and repeated sampling could better capture context-dependent bodily perturbations and their contribution to body-wandering. Third, some items differ in their level of specificity (e.g., heart or bladder versus the more general body), which may have influenced their rating distributions. This echoes the need for more detailed qualitative enquiry to better characterise these embodied thought dimensions.

Fourth, the scanner environment, while central to much of the literature on the neural correlates of mind-wandering, limits direct generalization to everyday settings (Linz et al., 2021; MacVittie et al., 2024; Wang et al., 2018). To address these constraints, future work should combine momentary experience sampling with experimental manipulations of interoception and arousal (Tyrer et al., 2025), and extend assessment to naturalistic contexts using ecological approaches (Linz et al., 2021; MacVittie et al., 2024). Indeed, consistent with our finding, a study implementing repetitive smartphone assessments in daily life also found that attention to interoceptive sensations were associated with negative emotional experience (Poerio et al., 2024). Finally, because both the content and metabolic demands of mind-wandering depend on task context and available energetic resources, it will be important to examine body-wandering across tasks varying in executive and affective demands, where embodied thought content and its coupling to psychophysiology may differ systematically (Birnie et al., 2015; Smallwood & Andrews-Hanna, 2013a).

### Conclusions

Together, these findings demonstrate the importance of accounting for the embodied contents of mind-wandering. Our results suggest that body-oriented thought represents a key feature of mind-wandering, co-varying with the contents, neurophysiological mechanisms, and affective implications of self-generated thought. They invite us to consider the stream of thought not only as a disengaged process of autobiographical projection, but also as one situated within the interoceptive context of the agent. Collectively, these findings support a view in which individual differences in the tendency to decouple from the present moment depends on both cognitive and interoceptive thought content, with body-wandering reflecting an engaged, affectively charged state anchored in the current bodily situation.

## Materials and methods

### Participants

We recruited 566 (360 females, 205 males, 1 other gender) participants (median age = 24, age range = 18-56) as part of the Visceral Mind Project, a large-scale neuroimaging study conducted at the Centre of Functionally Integrative Neuroscience, Aarhus University. We acquired participants via the SONA system participation pool at Aarhus University, or via local advertisements on posters and social media. Participants were provided with hourly compensation for their participation in the study. Participants had normal or corrected-to-normal vision and were fluent in Danish or English. Furthermore, we only included participants compatible with MRI scanning (e.g., not pregnant or breastfeeding, no metal implants, claustrophobia). Participants took part in multiple tasks, MRI scans, physiological recordings, and completed mental health and lifestyle inventories. In this study, we focus on the retrospective mind-wandering responses, as well as the resting state fMRI data, physiological recordings (photoplethysmography), and mental health assessments on depression and ADHD symptoms. The local Region Midtjylland Ethics Committee granted approval for the study and all participants provided informed consent. The study was conducted in accordance with the Declaration of Helsinki. 536 participants completed the post-resting state mind-wandering (338 females, 197 males, 1 other gender, median age = 24, age range = 18-56). After the removal of poor-quality fMRI data (see f/MRI preprocessing Methods for Quality Control procedures), a total of 489 full-dataset participants were included in the functional connectivity CCA analysis with body-wandering (311 females, 178 males, median age = 24, age range = 18-56) (see Supplementary Figure 16 for a full flow-chart of sample sizes).

### Multidimensional interoceptive experience sampling

We employed a multidimensional interoceptive experience sampling scale to retrospectively measure participants’ ongoing thoughts following the resting state scan. The scale consisted of an expanded version of the traditional multidimensional experiential sampling (MDES) technique by including items on interoceptive and somatic content (Martinon et al., 2019; Ruby et al., 2013; Smallwood et al., 2016, 2021; Wang et al., 2018), for a total of 22 items. Participants responded to all questions using a visual analogue scale, on a range of 0 to 100 (see Table 1). The presentation order of probe items was randomised across participants. The MDES probes were shown immediately after the 14-minute resting state fMRI scan with the following instructions: ‘Please answer a few questions about how you felt during the scan…’. They were previously unaware that they would be asked about their ongoing thoughts during the scan. We deliberately kept the instructions broad to keep the ratings not biased and open to the individual.

### Resting-state fMRI acquisition

We acquired anatomical and resting state fMRI data using a 3 T MRI scanner (Siemens Prisma) with a 32-channel head coil. We positioned small cushions around the head to minimise head movement. The participants wore earplugs and were instructed to keep as still as possible during the scans. The resting state scan included 600 volumes over 14 minutes using a T2*-weighted echo-planar imaging (EPI) multiband accelerated sequence (TR = 1400 ms, TE = 29.6 ms, voxel size = 1.79 x 1.79 x 1.80 mm). An acceleration factor of 4 was used in the slice direction along with GRAPPA in-plane acceleration factor of 2, and a flip-angle of 75°. A set of high-resolution whole brain T1-weighted anatomical images (0.9 mm^3 isotropic) were acquired using an MP-RAGE sequence (repetition time=2.2 s, echo time=2.51 ms, matrix size=256×256×192 voxels, flip angle=8°, AP acquisition direction). To increase comfort, all participants were instructed to use the bathroom just before scanning. They were instructed to have their eyes open during the scan, look at the fixation cross on a grey background, and not fall asleep.

### f/MRI Preprocessing

All fMRI data were preprocessed using fMRIPrep 22.1.1 (Esteban et al., 2019, 2022), built on Nipype 1.8.5 (Gorgolewski et al., 2018). The anatomical T1-weighted image was corrected for intensity non-uniformity using N4BiasFieldCorrection (Tustison et al., 2010) and ANTs 2.3.3, (Avants et al., 2008), skull-stripped, and segmented into gray matter, white matter, and cerebrospinal fluid using FAST (FSL 6.0.5.1) (Zhang et al., 2001). Brain surfaces were reconstructed using recon-all with FreeSurfer 7.2.0 (Dale et al., 1999), and the brain mask was estimated to reconcile ANTs-derived and FreeSurfer-derived cortical segmentations (Klein et al., 2017). Spatial normalization was performed using nonlinear registration to MNI152NLin2009cAsym and MNI152NLin6Asym templates via antsRegistration (ANTs 2.3.3).

For functional preprocessing, the resting-state fMRI were slice-time corrected using 3dTshift from AFNI ((Cox & Hyde, 1997), motion-corrected using MCFLIRT (FSL 6.0.5.1) (Jenkinson et al., 2002), and co-registered to the T1w reference using bbregister (FreeSurfer) (Greve & Fischl, 2009). Confounds including framewise displacement, DVARS, and CompCor components (combined anatomical and temporal) were computed (Behzadi et al., 2007; Jenkinson et al., 2002; Power et al., 2014). Motion estimates were expanded with temporal derivatives and quadratic terms. Functional data were resampled to native space and standard space (MNI152NLin2009cAsym) using a single interpolation step to minimize smoothing effects. Resampling to the fsaverage surface was also performed (Glasser et al., 2013). Poor quality fMRI data were excluded if any brain abnormalities were present, framewise displacement > 0.25, parts of brain image cut, unfinished recordings, or large motion artefacts in T1.

Functional connectivity analysis utilized the Schaefer atlas (200 cortical regions) with an additional 16 subcortical regions, and connectivity was estimated using Nilearn’s "ConnectivityMeasure" function, generating full 216-node correlation matrices. We low pass filtered at 0.08. Additional nuisance variable noise regression was performed using 24 motion parameters, noise signals from white matter and cerebral spinal fluid, six aCompCor ‘anat_combined’ parameters, and a high pass filter (Abraham et al., 2014; Behzadi et al., 2007).

For further details on the fMRI preprocessing workflow, please refer to the Supplementary Material.

### Physiological measurements

#### Physiological recording acquisition

We simultaneously recorded physiological measurements (photoplethysmography) during resting-state fMRI. The photoplethysmography was recorded on the right cheek via a Sentec Digital Monitoring system for the first data collection cohort at 1000 Hz (208 participants) and on the left index/middle finger via the MRI scanner for the second cohort at 200 Hz (325 participants). The first data collection cohort was recorded via a Brain Vision ExG system at 1000 Hz (208 participants), while the second cohort was recorded via an MRI scanner at 50 Hz (316 participants). Photoplethysmography recordings were acquired at 152.6 μV/bit resolution, and +/- 5000 mV range. See Supplementary Material for cardiac, respiratory, and gastric physiological data preprocessing (same protocol described in Banellis et al., 2025).

#### Cardiac preprocessing

R-peaks were identified from the cardiac recordings using the python package ‘systole’ using the ‘rolling_average_ppg’ method, while correcting for clipping artefacts in the second cohort’s data using a threshold of 950’000 and a cubic spline interpolation method. All the detected R-peaks were visually inspected and manually corrected, if necessary, as well as unintelligible segments marked and discarded. We excluded participants with less than 2 minutes of continuous cardiac recording. We computed heart rate as the mean of the beats per minute, while heart rate variability was defined as the root mean square of successive differences of the R-R intervals in milliseconds (RMSSD) (Legrand & Allen, 2022).

#### Physiological metric processing

We log transformed the RMSSD heart rate variability metric to ensure an approximate normal distributions. Furthermore, we removed outliers from all physiological metrics as 3 standard deviations from the mean. See Supplementary Material for further details of cardiac, respiratory, and gastric physiological data preprocessing (same protocol described in Banellis et al., 2025).

### Mental health assessment

We explored the mental health relationship of embodied and cognitive mind-wandering, specifically with individual differences in ADHD and depression symptoms. ADHD symptoms were assessed via the Adult ADHD Self-Report Scale (ASRS) using combined sum scores from both part A and part B (Kessler et al., 2005), while depression symptoms were examined via sum scores from the Major Depression Inventory (MDI) (with the highest score taken from 8a and 8b, 9a and 9b, 10a and 10b) (Bech et al., 2015). Psychological symptom assessment was completed online before attending the fMRI scan. We deliberately included participants ranging from subclinical symptom variation to symptom severity passing diagnostic clinical cut-offs. Specifically, 19% passed clinical thresholds for an ADHD diagnosis (binarised ASRS part A sum-score >=4), 2% for high levels of depression (MDI sum-score >30), 5% for moderate depression (MDI sum-score 26-30), and 6% for mild depression (MDI sum-score 21-25) (Kessler et al., 2005; Olsen et al., 2003). We log transformed the MDI sum scores to ensure an approximate normal distribution. Furthermore, we removed outliers from both the ASRS and MDI scores as 3 standard deviations from the mean.

### Data analysis

The mind-wandering item-level and mind-wandering average-level Wilcoxon signed-rank tests, partial correlations (controlling for age, gender, and BMI) were completed in JASP 0.18.3. The mind-wandering item-level cross correlations (corrected for multiple comparisons using the false discovery rate (FDR) using the Benjamini-Hochberg procedure at < 0.05 across the body-affect item correlations), mind-wandering item-level correlations with mental health symptoms and physiological metrics (corrected for multiple comparisons using the false discovery rate (FDR) using the Benjamini-Hochberg procedure at < 0.05 across the item correlations with mental health and physiology) was conducted in Rstudio 2023.03.0. Finally, the CCA of mind-wandering with functional connectivity and the gradient-based analysis was completed in Matlab 9.10.0.1710957 (R2021a), using the CCA-PLS toolbox (Mihalik et al., 2022).

#### Item cross correlation

We conducted exploratory Spearman’s cross-correlations on the body/mind-wandering item rating responses themselves, with p-values corrected for multiple comparisons using the false discovery rate. We also estimated partial correlations controlling for age, gender and body mass index.

#### Mind-wandering sampling validation

We assessed the construct validity of the mind-wandering items and examined their covariation via exploratory factor analysis (EFA). First, we inspected whether the mind-wandering responses were appropriate for factor analysis. We conducted Bartlett’s Test of Sphericity to ensure the correlation matrix of mind-wandering responses was not random (X2(210) = 1747.394, p < .001) (Bartlett, 1950). Furthermore, we ensured the Kaiser-Meyer-Olkin statistic was above 0.5 (MSA = 0.669) (Kaiser, 1974) to confirm factorability of the correlation matrix. Thus, the ‘Spontaneous’ item was removed from further analyses as the Kaiser-Meyer-Olkin statistic was < 0.5. For the EFA, we implemented a principal axis extraction method due to its robustness to nonnormality and sensitivity to identify weak constructs (Briggs & MacCallum, 2003). We assumed the mind-wandering items correlate, thus we employed an oblique (i.e., oblimin) rotation method. A Minimum Average Partial (MAP 2000) test indicated a two factor solution. Thus, we completed EFA on the mind-wandering responses inputting the following parameters: two factors, principal axis method and oblimin rotation. We completed the EFA analyses using the ‘fa’ function of the psych R statistical package. To evaluate the robustness of the factor solution, we conducted separate EFAs in even-row and odd-row participant subsets and calculated Pearson correlation coefficients between the resulting factor loadings. Finally, we assessed the relationship of outputted factor scores from the EFA with the mind-wandering summary scores via Spearman correlations.

#### Physiological and psychological fingerprints

We investigated the physiological and psychological fingerprints of mind-wandering items via false-discovery rate corrected Spearman’s correlation matrices. Additional partial correlations controlled for age, gender, and BMI.

To assess whether the observed psychophysiological associations generalised beyond individual mind-wandering items, we repeated the mental health and psychophysiological correlation analyses using latent variables derived from the exploratory factor analysis (Figure 3) and the canonical correlation analysis (Figure 4). Specifically, Spearman correlations were computed between (i) EFA-derived factor scores and (ii) CCA-derived variates with mental health symptom scores and physiological metrics.

#### Canonical correlation analysis

In this study, we used the CCA-PLS toolbox to fit cross-validated Canonical Correlation Analysis (CCA) models relating individual whole-brain functional connectivity matrices to individual body-wandering and cognitive-wandering (MDES) items. The CCA-PLS toolbox is a MATLAB-based software designed for investigating multivariate associations between multiple modalities of data, such as brain imaging and behavioural measures (Mihalik et al., 2022). Specifically, CCA aims to find linear combinations of each multidimensional variable (i.e., canonical variates, which are weighted sums: connectivity (i.e., V = X * B) and body/cognitive-wandering (i.e., U = Y * A)) that are maximally correlated with each other, but uncorrelated with all other combinations (X and Y represent the inputted data, while A and B represent the canonical weights/vectors) (Linke et al., 2021; Mihalik et al., 2022). The toolbox incorporates various CCA/PLS models, including the cross-validated and optimised PCA-CCA techniques applied here.

We began by preprocessing the whole-brain functional connectivity matrices and individual MDES items, ensuring that each dataset was standardised to have zero mean and unit variance. Furthermore, we regressed nuisance covariates (i.e., gender, age, BMI, data collection cohort) from both the connectivity and MDES item variables. Subsequently, we applied the cross-validated CCA approach within the predictive framework (i.e., machine learning) provided by the CCA-PLS toolbox. This predictive approach involved randomly splitting the data into an optimization/training set (i.e., 60% of the overall data) and a holdout/test set (i.e., 40% of the overall data) 5 times (i.e., 5-fold cross-validation). These optimization and holdout sets are known as the ‘outer data splits’, used for statistical inference (i.e., determining the number of significant associative CCA effects/modes). The p-values were calculated via permutation testing (i.e., 1000 permutations), as the fraction of the shuffled permuted out-of-sample correlations exceeding the out-of-sample correlation in the non-permuted holdout set. Because we implemented 5 holdout sets, the p-value for each holdout set was Bonferroni corrected (i.e., *a* = 0.05/5 = 0.01). An associative CCA effect was considered significant if the p-value was significant in at least one of the holdout sets. If a significant associative CCA effect was found, the CCA iteratively removes the effect from the data via deflation and repeats this approach to find orthogonal CCA associative effects.

Furthermore, because we conducted PCA before CCA (to reduce the connectivity data and body/cognitive mind-wandering item data dimensions), we further divided the optimization set into a training set (i.e., 60% of the optimization set) and a validation set (i.e., 40% of the optimization set) 5 times (i.e., 5-fold hyperparameter optimization). These ‘inner data splits’ were used to select the optimal hyperparameters (i.e., number of PCA components for each dataset) by maximising the average out-of-sample correlation in the validation sets (i.e., model generalizability). In addition to evaluating model generalizability via the predictive approach, we assessed the stability of the CCA models by measuring the average similarity of weights across different training sets. Furthermore, we assessed the robustness (i.e., number of significant data splits) of the CCA model (80%).

To visualise the results, we plotted the maximally correlated canonical variate via a scatter plot, and the connectivity and body/cognitive-wandering structure correlations (i.e., Pearson’s correlations between inputted variables and respective canonical variates) using a chord plot, and bar plot/word cloud, respectively. Note, weights were back-projected from PCA feature space to input space to facilitate the relationships with the original input variables. Bar plots and word clouds were used to display the individual mind-wandering item loadings (structure correlations of raw items with the mind-wandering variate), while chord plots provided a graphical representation of the connections between brain regions and their corresponding loadings (structure correlations of raw functional connectivity data with the functional connectivity variate), thresholded at showing the top 99th percentile of the connectivity loadings. To visualise the brain networks contributing to the CCA result, we organised top connectivity loadings in the chord plot according to yeo-7 networks (Yeo et al., 2011). In addition, recently there has been interest in the properties of an ‘interoceptive-allostatic network’ (Kleckner et al., 2017). To define this network we reviewed the neuroanatomy and functional imaging literature on interoception and defined cortical regions in the interoceptive network as follows: insular cortex, cingulate cortex, frontal orbital cortex, frontal medial cortex, postcentral gyrus (primary somatosensory cortex), and central opercular cortex (secondary somatosensory cortex) (Azzalini et al., 2019; Barrett & Simmons, 2015; Craig, 2002; Engelen et al., 2023; Khalsa et al., 2009; Kleckner et al., 2017). Furthermore, we illustrated the highest contributing nodes by summing the amount of edge connections above the 99th percentile cutoff for each node. This was plotted on a standard mni152 brain template using MRIcroGL at the following MNI coordinates: 8, −20, 28. These visualisations allowed for a clearer understanding of the associations between the whole-brain functional connectivity matrices and individual body/cognitive-wandering items, highlighting the key brain-behaviour relationships identified by our cross-validated CCA model.

#### Gradient map association

We investigated the localisation of the body-wandering CCA result in relation to five brain variation gradient maps from the Human Connectome Project resting connectivity data (Margulies et al., 2016). These gradients (five cortical and subcortical surface atlases) were extracted from Neurovault (https://neurovault.org/collections/1598/). The combined 200-cortical-region Schaefer-16-subcortical-region atlas was converted to surface space using the freesurfer function ‘*mri_vol2surf’* (this removed the bilateral caudate as these deep subcortical regions were not represented in surface space). For each of the five gradient maps, we averaged the gradient values of the top CCA brain parcels (parcels which passed the top 1% threshold of the connectivity CCA loadings) and generated a null distribution with 2500 permutations via a spin-test which shuffles the gradient values. The p-values were generated by comparing the overall gradient average across the top CCA brain regions, with the 2500 null averages generated from the spin-test. The five p-values computed from repeating this process for each of the five gradient atlases were Bonferroni corrected (i.e., *a* = 0.05/5 = 0.01).

#### Convergent validity of the body-wandering connectome

To assess the convergent validity of the embodied dimension of thought across analytical approaches, we tested whether the brain connectivity pattern associated with the unsupervised behavioural factor derived from exploratory factor analysis (EFA) overlapped with the connectivity profile identified by the canonical correlation analysis (CCA). For each participant, whole-brain functional connectivity matrices were vectorised (upper triangle, excluding the diagonal). A mass-univariate regression was then performed across all connectivity edges (n = 23,220), relating individual EFA factor scores to edge-wise connectivity strength using ordinary least squares with non-parametric permutation testing (10,000 permutations), as implemented in nilearn’s ‘permuted_ols’ function. This procedure was conducted separately for Factor 1 (Body-Wandering) and Factor 2 (Mental Time Travel), yielding edge-wise regression coefficients and permutation-derived p-values.

To quantify overlap between the unsupervised EFA-derived connectivity pattern and the supervised CCA solution, we computed Spearman correlations between the vector of mass-univariate connectivity coefficients and the CCA brain connectivity loadings across all edges. Statistical significance of these correlations was assessed using permutation testing (10,000 permutations).

#### Control analyses

To ensure that our findings were not driven by individual differences in physical discomfort or affective traits, we conducted a series of control analyses accounting for state-based bladder-related discomfort, trait-level somatic, and trait anxiety. These controls were applied consistently across both the analyses involving psychophysiological correlations with mind-wandering items, and the CCA of mind-wandering items with brain connectivity. For the psychophysiological correlation analysis, this involved regressing individuals scores on the control variable from the mind-wandering items, before conducting the correlations with psychophysiological variables. For the CCA with brain connectivity, we included the control variable as a nuisance regressor with age, gender, and BMI. All other parameters of the CCA were identical to the analysis described above. This procedure was performed separately for momentary bladder sensation (using individual ratings for the ‘Bladder’ item), trait-level somatic distress (using the Patient Health Questionnaire somatic symptoms module, PHQ-15 (Kroenke et al., 2002)), and trait anxiety (using the State-Trait Anxiety Inventory, STAI-T (Spielberger, 1983)).

#### Post-hoc power analyses

We did a post-hoc power analysis based on the effect size from the CCA result of mind-wandering with brain connectivity. The canonical correlation coefficient from the CCA was r = 0.354, which was converted to an effect size of f^2^ = 0.14327. To determine the required sample size for 80% power at an alpha level of 0.05, we conducted the analysis using the ‘pwr.f2.test’ function in R. The number of variables included was 17 (2 brain connectivity, 15 mind-wandering items, PCA-reduced for the reported data-fold), with 134 degrees of freedom. Based on this analysis, we determined that a minimum sample size of 152 participants was required to achieve adequate power for detecting the observed effect in the CCA model. Our actual CCA sample size of 489 is approximately 3.22 times larger than the estimated required sample size, providing substantially greater statistical power.

## Data availability

Deidentified participant data and scripts implemented in this paper are available here: https://github.com/embodied-computation-group/body_wandering_CCA

## Acknowledgements

This research is financially supported by a Lundbeckfonden Fellowship (R272-2017-4345) and a European Research Council Grant (ERC-2020-StG-948788) awarded to MGA. FF is supported by a European Research Council Grant (ERC-2020-StG-948838) and a Lundbeckfonden Experiment Grant (R436-2023-991). The funding sources were not involved in the study design, collection, analysis, interpretation, or writing of the manuscript.

We dedicate this paper to the memory of Jonny Smallwood, a coauthor and close friend. His ideas, intellectual generosity, and commitment to careful theory deeply shaped this work and continue to influence our thinking.

## Author contributions

LB analysed the data, interpreted the results, and wrote the manuscript, NN collected the data, provided conceptual advice and contributed towards preprocessing of neuroimaging data, MB collected the data and contributed towards the preprocessing of the physiological data, MN collected the data, IG provided conceptual advice on physiological data acquisition, and contributed towards the preprocessing of the physiological data, NL collected the data, and contributed towards preprocessing, FF provided conceptual advice and edited the manuscript, JS provided conceptual advice, analysis supervision, and wrote the manuscript, MA provided supervision, conceptual advice, and wrote the manuscript.

## Materials and correspondence

Leah Banellis.

