## Supplementary Material for "Uncovering the embodied dimension of the wandering mind"

**Supplementary Table 1:** Wilcoxon signed-rank tests of movement/body vs interoceptive mind-wandering items.

| Measure 1 | | Measure 2 | W | z | *p* | Rank-Biserial Correlation | SE Rank-Biserial Correlation |
|---|---|---|---|---|---|---|---|
| Movement | - | Breath | 69170 | 0.343 | 0.732 | 0.017 | 0.051 |
| Movement | - | Heart | 75063 | 3.062 | 0.002 | 0.157 | 0.051 |
| Movement | - | Bladder | 93763 | 10.171 | < .001 | 0.528 | 0.052 |
| Movement | - | Skin | 91493.5 | 6.932 | < .001 | 0.351 | 0.051 |
| Movement | - | Stomach | 80078 | 4.663 | < .001 | 0.239 | 0.051 |
| Body | - | Breath | 78066 | 3.878 | < .001 | 0.198 | 0.051 |
| Body | - | Heart | 88206 | 7.021 | < .001 | 0.359 | 0.051 |
| Body | - | Bladder | 101878 | 12.836 | < .001 | 0.667 | 0.052 |
| Body | - | Skin | 97973 | 9.752 | < .001 | 0.498 | 0.051 |
| Body | - | Stomach | 93243.5 | 8.234 | < .001 | 0.420 | 0.051 |

**Supplementary Table 2:** Spearmen correlations of body-wandering items with affective mind-wandering items (significant in bold, negative correlations in blue, and positive correlations in red).

| | Pos | Neg |
|---|---|---|
| Arousal | **rs = -0.175, *p*FDR = <.001** | rs = 0.013, *p*FDR = .796 |
| Body | rs = -0.035, *p*FDR = .471 | **rs = 0.118, *p*FDR = .012** |
| Breath | rs = -0.042, *p*FDR = .398 | **rs = 0.173, *p*FDR = <.001** |
| Heart | **rs = -0.267, *p*FDR = <.001** | **rs = 0.155, *p*FDR = .001** |
| Movement | **rs = 0.102, pFDR = .030** | rs = -0.009., *p*FDR = .837 |
| Bladder | **rs = -0.298, *p*FDR = <.001** | **rs = 0.186, *p*FDR = <.001** |
| Skin | **rs = -0.287, *p*FDR = <.001** | **rs = 0.225, *p*FDR = <.001** |
| Stomach | **rs = -0.183, *p*FDR = <.001** | **rs = 0.163, *p*FDR = <.001** |

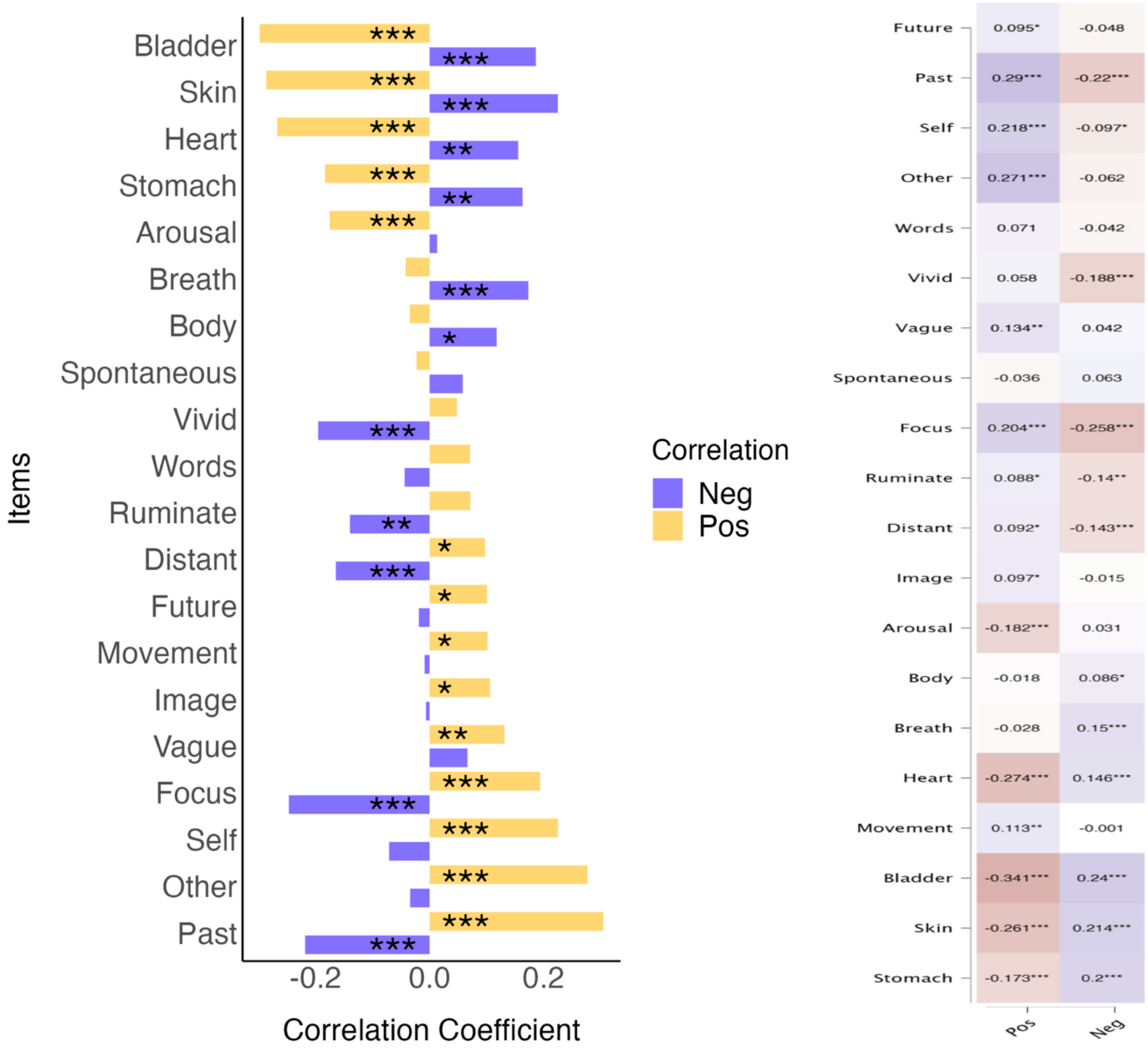


**Supplementary Figure 1:** (Left) All mind-wandering item Spearman correlations with the affective items (Pos and Neg) and (Right) their partial Spearman correlations (controlled for age, gender, BMI). Asterisks indicate the significance of the correlations in both plots (* $p \leq 0.05$, ** $p \leq 0.01$, ***$p \leq 0.001$).

**Supplementary Table 3**: Spearman correlations of mind-wandering items with mental health (ADHD and depression) scores and physiological metrics of arousal (heart rate and heart rate variability RMSSD) (significant in bold, negative correlations in blue, and positive correlations in red).

| | ADHD (ASRS) | Depression (MDI) | Heart Rate | HRV RMSSD |
|---|---|---|---|---|
| Future | $r_s$ = 0.074, pFDR = .173 | **$r_s$ = 0.123, pFDR = .016** | $r_s$ = -0.057, pFDR = .303 | **$r_s$ = 0.133, pFDR = .008** |
| Past | $r_s$ = 0.090, pFDR = .087 | **$r_s$ = 0.164, pFDR < .001** | $r_s$ = -0.095, pFDR = .064 | **$r_s$ = 0.138, pFDR = .006** |
| Self | $r_s$ = 0.099, pFDR = .055 | $r_s$ = 0.046, pFDR = .409 | $r_s$ = -0.001, pFDR = .974 | $r_s$ = 0.096, pFDR = .063 |
| Other | $r_s$ = 0.082, pFDR = .121 | $r_s$ = 0.046, pFDR = .411 | $r_s$ = -0.037, pFDR = .510 | **$r_s$ = 0.135, pFDR = .007** |
| Pos | $r_s$ = 0.086, pFDR = .102 | $r_s$ = 0.094, pFDR = .072 | $r_s$ = -0.068, pFDR = .202 | **$r_s$ = 0.157, pFDR = .001** |
| Neg | $r_s$ = 0.010, pFDR = .870 | $r_s$ = -0.047, pFDR = .409 | **$r_s$ = 0.120, pFDR = .018** | **$r_s$ = -0.140, pFDR = .005** |
| Words | $r_s$ = 0.036, pFDR = .523 | $r_s$ = -0.042, pFDR = .450 | $r_s$ = 0.037, pFDR = .510 | $r_s$ = 0.065, pFDR = .234 |
| Vivid | $r_s$ = -0.082, pFDR = .124 | $r_s$ = -0.025, pFDR = .670 | $r_s$ = -0.020, pFDR = .737 | $r_s$ = 0.026, pFDR = .651 |
| Vague | $r_s$ = 0.096, pFDR = .063 | $r_s$ = 0.086, pFDR = .104 | $r_s$ = -0.086, pFDR = .101 | $r_s$ = 0.090, pFDR = .084 |
| Spontan | $r_s$ = 0.013, pFDR = .833 | $r_s$ = -0.063, pFDR = .250 | $r_s$ = -0.004, pFDR = .946 | $r_s$ = -0.011, pFDR = .855 |
| Focus | $r_s$ = 0.023, pFDR = .686 | $r_s$ = 0.074, pFDR = .174 | $r_s$ = -0.035, pFDR = .526 | **$r_s$ = 0.130, pFDR = .009** |
| Ruminate | $r_s$ = 0.069, pFDR = .200 | $r_s$ = 0.012, pFDR = .845 | $r_s$ = -0.043, pFDR = .433 | $r_s$ = 0.071, pFDR = .190 |
| Distant | $r_s$ = -0.046, pFDR = .409 | $r_s$ = 0.002, pFDR = .961 | $r_s$ = 0.024, pFDR = .674 | $r_s$ = 0.014, pFDR = .824 |
| Image | $r_s$ = 0.034, pFDR = .538 | $r_s$ = 0.099, pFDR = .055 | $r_s$ = -0.029, pFDR = .615 | $r_s$ = 0.023, pFDR = .695 |
| Arousal | $r_s$ = -0.089, pFDR = .089 | **$r_s$ = -0.108, pFDR = .036** | $r_s$ = 0.024, pFDR = .674 | $r_s$ = -0.049, pFDR = .379 |
| Body | $r_s$ = 0.019, pFDR = .742 | $r_s$ = -0.046, pFDR = .409 | $r_s$ = 0.014, pFDR = .824 | $r_s$ = -0.020, pFDR = .729 |
| Breath | $r_s$ = -0.004, pFDR = .946 | $r_s$ = 0.059, pFDR = .281 | $r_s$ = 0.065, pFDR = .224 | $r_s$ = -0.051, pFDR = .366 |
| Heart | **$r_s$ = -0.211, pFDR < .001** | **$r_s$ = -0.149, pFDR = .003** | $r_s$ = 0.009, pFDR = .875 | $r_s$ = -0.046, pFDR = .409 |
| Movement | $r_s$ = 0.068, pFDR = .206 | $r_s$ = 0.039, pFDR = .491 | $r_s$ = -0.006, pFDR = .912 | $r_s$ = 0.019, pFDR = .742 |
| Bladder | **$r_s$ = -0.106, pFDR = .040** | **$r_s$ = -0.133, pFDR = .008** | **$r_s$ = 0.119, pFDR = .019** | **$r_s$ = -0.149, pFDR = .003** |
| Skin | **$r_s$ = -0.104, pFDR = .046** | **$r_s$ = -0.129, pFDR = .011** | **$r_s$ = 0.150, pFDR = .003** | **$r_s$ = -0.234, pFDR < .001** |
| Stomach | $r_s$ = -0.073, pFDR = .180 | **$r_s$ = -0.105, pFDR = .043** | $r_s$ = 0.090, pFDR = .086 | $r_s$ = -0.078, pFDR = .145 |

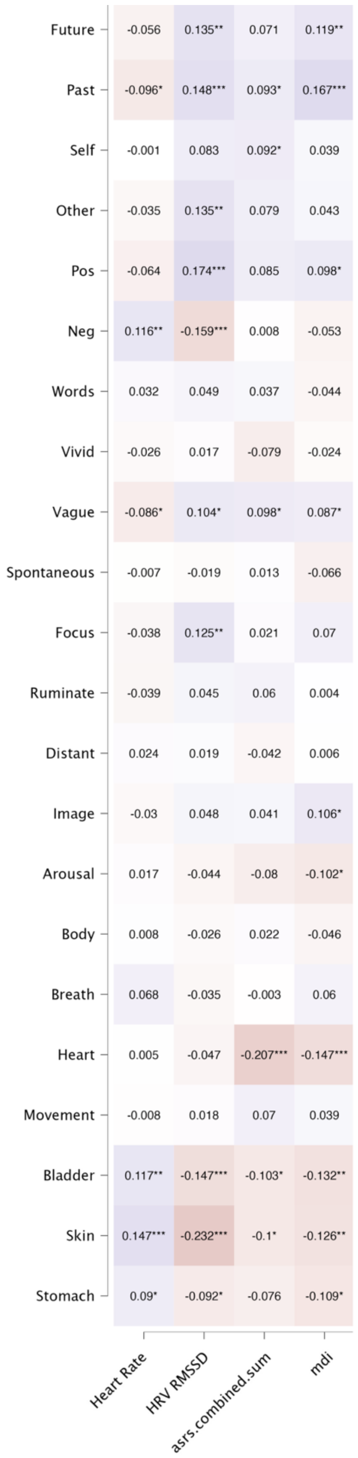


**Supplementary Figure 2:** Mind-wandering item partial Spearman correlations with psychological and physiological variables (controlled for age, gender, BMI). Asterisks indicate the significance of the partial correlation (* $p \leq 0.05$, ** $p \leq 0.01$, ***$p \leq 0.001$), while in red are negative correlations and in blue are positive correlations.

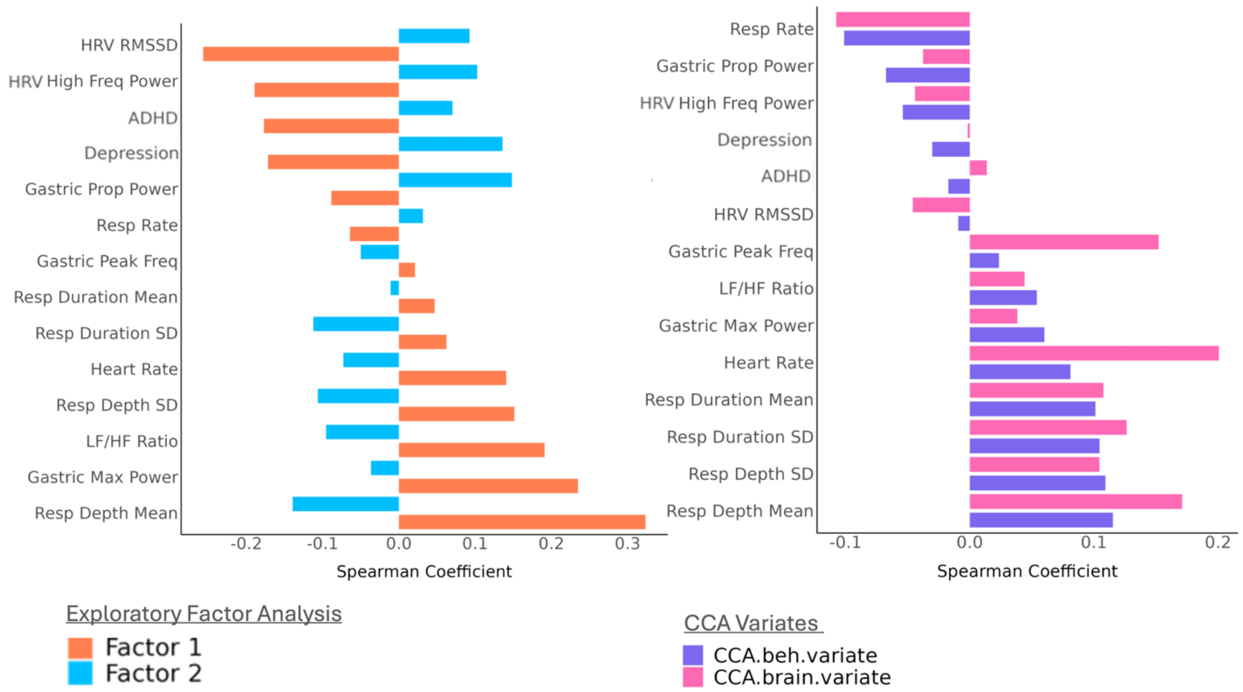


**Supplementary Figure 3:** Psychophysiological Spearman correlations with (Left) factor scores from the exploratory factor analysis (Figure 3 in main manuscript) and (Right) variates from the canonical correlation analysis (Figure 4 in main manuscript). This demonstrates a similar anticorrelated pattern to the body-wandering and cognitive/descriptive mind-wandering item correlations with psychophysiology showed in Figure 2.

**Supplementary Table 4:** The items of the mental health surveys (Adult ADHD Self-Report Scale and Major Depression Inventory).

| Item | Question | Response |
|---|---|---|
| | **Adult ADHD Self-Report Scale (ASRS)** | |
| asrs_a_1 | How often do you have trouble wrapping up the final details of a project, once the challenging parts have been done? | 1 (Never) – 5 (Very Often) |
| asrs_a_2 | How often do you have difficulty getting things in order when you have to do a task that requires organization? | 1 (Never) – 5 (Very Often) |
| asrs_a_3 | How often do you have problems remembering appointments or obligations? | 1 (Never) – 5 (Very Often) |
| asrs_a_4 | When you have a task that requires a lot of thought, how often do you avoid or delay getting started? | 1 (Never) – 5 (Very Often) |
| asrs_a_5 | How often do you fidget or squirm with your hands or feet when you have to sit down for a long time? | 1 (Never) – 5 (Very Often) |
| asrs_a_6 | How often do you feel overly active and compelled to do things, like you were driven by a motor? | 1 (Never) – 5 (Very Often) |
| asrs_b_1 | How often do you make careless mistakes when you have to work on a boring or difficult project? | 1 (Never) – 5 (Very Often) |
| asrs_b_2 | How often do you have difficulty keeping your attention when you are doing boring or repetitive work? | 1 (Never) – 5 (Very Often) |
| asrs_b_3 | How often do you have difficulty concentrating on what people say to you, even when they are speaking to you directly? | 1 (Never) – 5 (Very Often) |
| asrs_b_4 | How often do you misplace or have difficulty finding things at home or at work? | 1 (Never) – 5 (Very Often) |
| asrs_b_5 | How often are you distracted by activity or noise around you? | 1 (Never) – 5 (Very Often) |
| asrs_b_6 | How often do you leave your seat in meetings or other situations in which you are expected to remain seated? | 1 (Never) – 5 (Very Often) |
| asrs_b_7 | How often do you feel restless or fidgety? | 1 (Never) – 5 (Very Often) |
| asrs_b_8 | How often do you have difficulty unwinding and relaxing when you have time to yourself? | 1 (Never) – 5 (Very Often) |
| asrs_b_9 | How often do you find yourself talking too much when you are in social situations? | 1 (Never) – 5 (Very Often) |

| | | |
|---|---|---|
| asrs_b_10 | When you're in a conversation, how often do you find yourself finishing the sentences of the people you are talking to, before they can finish them themselves? | 1 (Never) – 5 (Very Often) |
| asrs_b_11 | How often do you have difficulty waiting your turn in situations when turn taking is required? | 1 (Never) – 5 (Very Often) |
| asrs_b_12 | How often do you interrupt others when they are busy? | 1 (Never) – 5 (Very Often) |

**Major Depression Inventory (MDI)**

| | | |
|---|---|---|
| mdi_1 | Have you felt low in spirits or sad? | 0 (At no time) – 5 (All the time) |
| mdi_2 | Have you lost interest in your daily activities? | 0 (At no time) – 5 (All the time) |
| mdi_3 | Have you felt lacking in energy and strength? | 0 (At no time) – 5 (All the time) |
| mdi_4 | Have you felt less self-confident? | 0 (At no time) – 5 (All the time) |
| mdi_5 | Have you had a bad conscience or feelings of guilt? | 0 (At no time) – 5 (All the time) |
| mdi_6 | Have you felt that life wasn't worth living? | 0 (At no time) – 5 (All the time) |
| mdi_7 | Have you had difficulty in concentrating, e.g. when reading the newspaper or watching TV? | 0 (At no time) – 5 (All the time) |
| mdi_8a | Have you felt very restless? | 0 (At no time) – 5 (All the time) |
| mdi_8b | Have you felt subdued or slowed down? | 0 (At no time) – 5 (All the time) |
| mdi_9a | Have you been sleeping too little? | 0 (At no time) – 5 (All the time) |
| mdi_9b | Have you been sleeping too much? | 0 (At no time) – 5 (All the time) |
| mdi_10a | Have you suffered from reduced appetite? | 0 (At no time) – 5 (All the time) |
| mdi_10b | Have you suffered from increased appetite? | 0 (At no time) – 5 (All the time) |

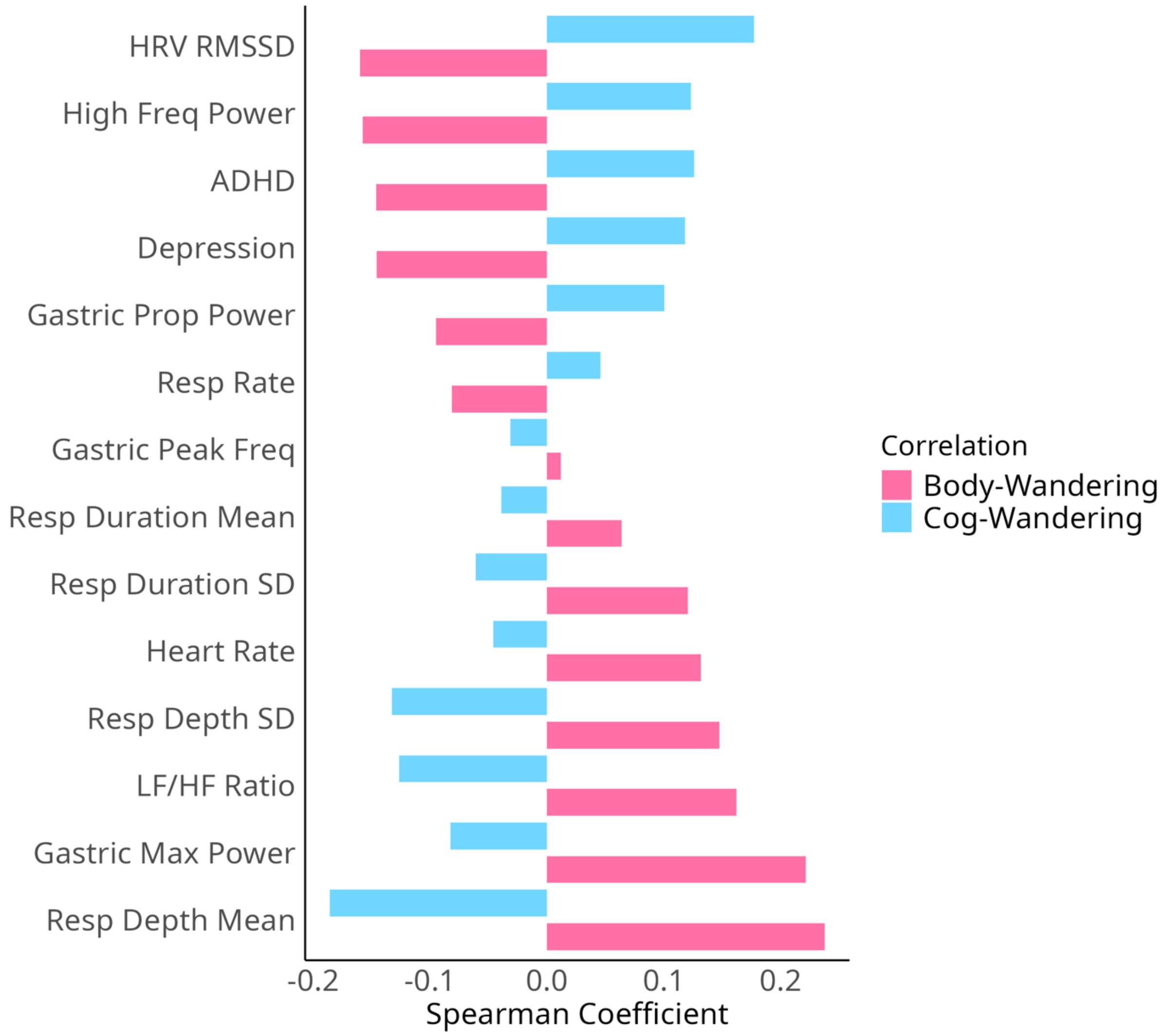


**Supplementary Figure 4:** Mind-wandering Spearman correlations with psychological (ADHD and depression scores) and physiological variables (across cardiac, respiratory, and gastric metrics). In blue are correlations using an average of the 14 original mind-wandering items and in pink are correlations with an average of the additional 8 body-related items (see Figure 1 and Table 1 in main manuscript for item details). This demonstrates an anticorrelated pattern of body-wandering relations with psychophysiological variables, in comparison to descriptive/cognitive-wandering correlations with psychophysiology.

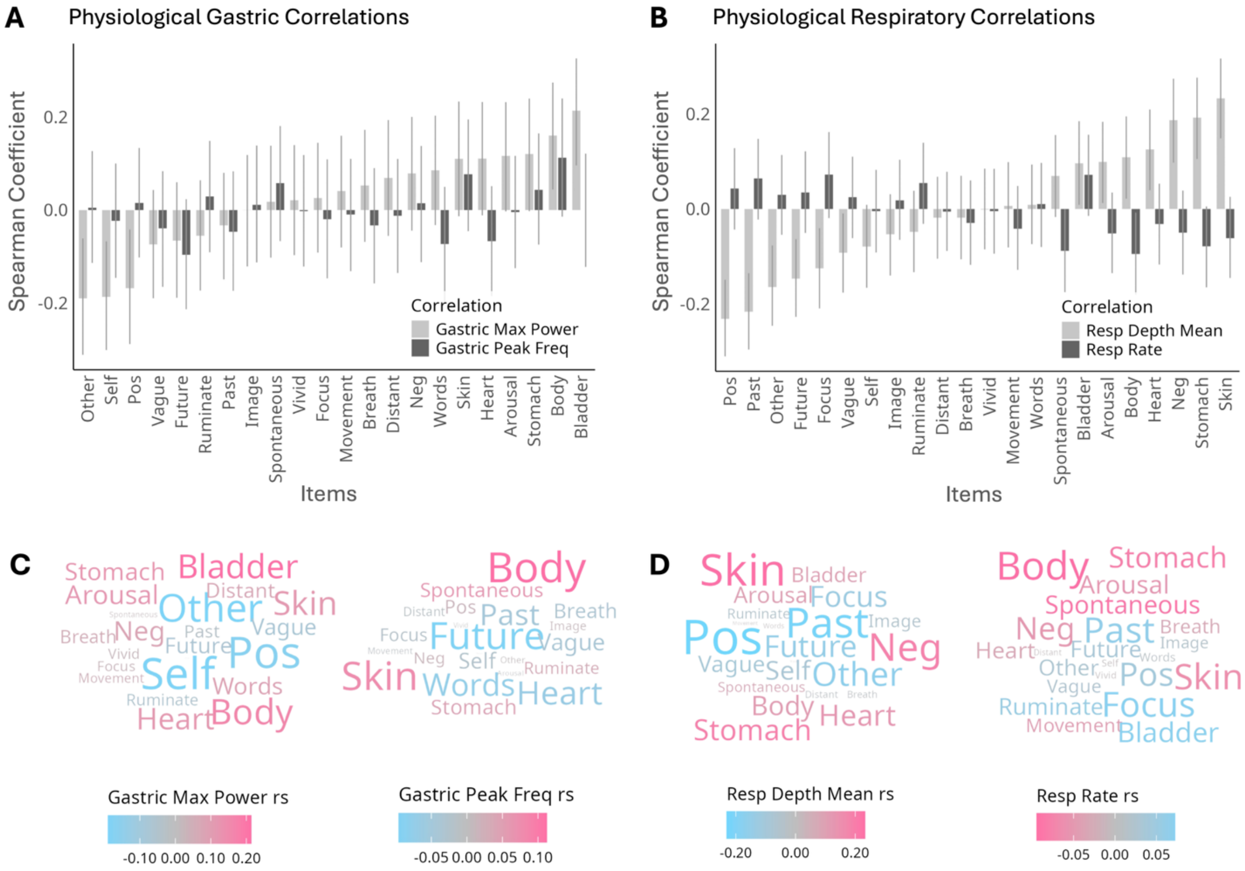


**Supplementary Figure 5:** Mind-wandering item Spearman correlations with gastric and respiratory physiological variables. (A and C) Gastric metric correlations revealing higher gastric maximum power with somatic items, whereas social-cognitive are associated with reduced gastric power. (B and D) In the respiratory domain, somatic attention is characterized by deeper respiratory breaths, while cognitive wandering reflects a shallower respiratory breath.

EFA Loadings (Thresholded at |0.3|)

| Item | Factor 1 | Factor 2 |
|---|---|---|
| Future | -0.06 | 0.3 |
| Past | -0.24 | **0.4** |
| Self | **-0.35** | -0.11 |
| Other | **-0.39** | 0.06 |
| Pos | **-0.73** | 0.04 |
| Neg | **0.53** | 0.03 |
| Words | -0.1 | -0.17 |
| Vivid | 0.05 | 0.19 |
| Vague | -0.01 | **0.64** |
| Focus | -0.08 | **0.35** |
| Ruminate | -0.22 | **-0.35** |
| Distant | -0.03 | 0.15 |
| Image | 0.03 | **0.62** |
| Arousal | 0.28 | 0.07 |
| Body | 0.1 | -0.13 |
| Breath | 0.14 | 0.22 |
| Heart | **0.36** | -0.15 |
| Movement | -0.03 | -0.06 |
| Bladder | **0.35** | -0.27 |
| Skin | **0.48** | 0.16 |
| Stomach | **0.31** | 0.07 |

**Supplementary Figure 6:** Exploratory Factor Analysis of the Multidimensional Interoceptive Experience Sampling data, revealing two factors. This shows the loadings (pattern coefficients) with bold representing > absolute 0.3.

**Supplementary Table 5:** ***Cross-validated CCA signifying functional connectivity fingerprint of body-wandering.***

| | **Connectivity** | | **Mind-Wandering** | | **Relationship** | | |
|---|---|---|---|---|---|---|---|
| | **Weight Stability** | **Explained Variance** | **Weight Stability** | **Explained Variance** | **In-Sample Correlation** | **Out-of-Sample Correlation** | **Robust-ness** |
| **CCA Mode** | 0.953 | 4.529 | 0.642 | 5.419 | 0.386 | 0.354 | 80% |

The CCA mode characteristics which define the functional connectivity fingerprint of body-wandering. This includes the ‘weight stability’ describing the stability of the CCA model via the average similarity (Pearson’s correlation) of weights across different training sets of the outer data splits. The percent ‘explained variance’ of the CCA model in contrast to the variance across the training sets of the outer data splits. The ‘in-sample correlation’ of the canonical variates between the training sets of the outer data splits. The ‘out-of-sample correlation’ of the canonical variates between the test sets of the outer data splits (cross-validated CCA relationship). Robustness indicates that 4/5 of the data folds (80% of the cross-validated iterations) revealed a significant relationship between the training and test set, and therefore we demonstrate stability and generalisability of the functional connectivity-body wandering signature across the data folds.

**Supplementary Table 6:** Highest connectivity CCA loadings for the top 35 brain connectivity edges. The region A/B cortical labels are from the Harvard-Oxford Cortical Structural Atlas, while the network A/B labels are from the Yeo-7 network assignment (with the regions relabelled to 'Intero' for visualisation shown in brackets).

| Region A | Region B | Canonical Loading | Network A | Network B |
|---|---|---|---|---|
| Precentral.Gyrus.R.1 | Thalamus.R | 0.703031348390156 | SomMot | Subcortical |
| Precentral.Gyrus.L.2 | Caudate.R | 0.698837570190837 | SomMot | Subcortical |
| Precentral.Gyrus.R.2 | Thalamus.R | 0.695841268698783 | SomMot | Subcortical |
| Postcentral.Gyrus.R.1 | Thalamus.R | 0.694528867386115 | SomMot (Intero) | Subcortical |
| Precentral.Gyrus.L.1 | Thalamus.R | 0.683341470686942 | SomMot | Subcortical |
| Postcentral.Gyrus.R | Thalamus.R | 0.681264971081168 | SomMot (Intero) | Subcortical |
| Postcentral.Gyrus.R.3 | Paracingulate.Gyrus.R | 0.678690482256079 | SomMot (Intero) | Cont (Intero) |
| Postcentral.Gyrus.R.4 | Caudate.R | 0.675770903449253 | DorsAttn (Intero) | Subcortical |
| Precentral.Gyrus.L.4 | Caudate.R | 0.674086239587976 | SomMot | Subcortical |
| Juxtapositional.Lobule.Cortex..L | Thalamus.R | 0.672817793626145 | SomMot | Subcortical |
| Postcentral.Gyrus.L.1 | Caudate.R | 0.672462998807524 | SomMot (Intero) | Subcortical |
| Precentral.Gyrus.L.3 | Thalamus.R | 0.669099993214319 | SomMot | Subcortical |
| Parietal.Operculum.Cortex.R | Thalamus.R | 0.668969113143266 | SomMot | Subcortical |
| Precentral.Gyrus.L | Thalamus.R | 0.668247273889767 | SomMot | Subcortical |
| Postcentral.Gyrus.R.5 | Caudate.R | 0.667229739152148 | DorsAttn (Intero) | Subcortical |
| Postcentral.Gyrus.L.2 | Caudate.R | 0.666611048372896 | SomMot (Intero) | Subcortical |
| Postcentral.Gyrus.R.2 | Caudate.R | 0.666145278761403 | SomMot (Intero) | Subcortical |
| Juxtapositional.Lobule.Cortex..R | Thalamus.R | 0.666016991378321 | SomMot | Subcortical |
| Planum.Temporale.R | Thalamus.R | 0.665639737469437 | SomMot | Subcortical |
| Postcentral.Gyrus.R.3 | Caudate.R | 0.664555895370312 | SomMot (Intero) | Subcortical |
| Superior.Frontal.Gyrus.R | Thalamus.R | 0.660599005722293 | SomMot | Subcortical |
| Precentral.Gyrus.L.4 | Putamen.L | 0.66007848150924 | SomMot | Subcortical |

| | | | | |
|---|---|---|---|---|
| Postcentral.Gyrus.L.1 | Paracingulate.Gyrus.R | 0.660041933513838 | SomMot (Intero) | Cont (Intero) |
| Superior.Parietal.Lobule.R | Thalamus.R | 0.659477978020345 | SomMot | Subcortical |
| Postcentral.Gyrus.R.2 | Caudate.L | 0.658217710091078 | SomMot (Intero) | Subcortical |
| Postcentral.Gyrus.L.3 | Thalamus.R | 0.658135150734579 | SomMot (Intero) | Subcortical |
| Postcentral.Gyrus.R.3 | Thalamus.R | 0.657372431563344 | SomMot (Intero) | Subcortical |
| Insular.Cortex.R | Thalamus.R | 0.656215782982937 | DorsAttn (Intero) | Subcortical |
| Precentral.Gyrus.L.2 | Putamen.L | 0.654595727561157 | SomMot | Subcortical |
| Postcentral.Gyrus.R.6 | Thalamus.R | 0.65228900504608 | Cont (Intero) | Subcortical |
| Precentral.Gyrus.L.4 | Thalamus.R | 0.650566303406784 | SomMot | Subcortical |
| Postcentral.Gyrus.L.1 | Thalamus.R | 0.650524205652741 | SomMot (Intero) | Subcortical |
| Central.Opercular.Cortex.R | Thalamus.R | 0.650409457802497 | SomMot | Subcortical |
| Postcentral.Gyrus.L | Caudate.R | 0.649485513639986 | Default (Intero) | Subcortical |
| Postcentral.Gyrus.R.1 | Paracingulate.Gyrus.R | 0.648643551324168 | SomMot (Intero) | Cont (Intero) |

**Supplementary Table 7:** The summed total number of edges for each of the top 1% (> absolute 0.558) connectivity CCA loading brain regions (i.e., edge sums or connectivity degree), and their respective network. The region labels are from the Harvard-Oxford Cortical Structural Atlas, while the network labels are from the Yeo-7 network assignment (with the regions relabelled to 'Intero' for visualisation shown in brackets).

| Region | Network | Degree (Edge Sums) |
|---|---|---|
| Thalamus R | Subcortical | 53 |
| Postcentral Gyrus R | SomMot (Intero) | 24 |
| Thalamus L | Subcortical | 22 |
| Caudate R | Subcortical | 22 |
| Putamen L | Subcortical | 21 |
| Postcentral Gyrus L | SomMot (Intero) | 18 |
| Postcentral Gyrus R | SomMot (Intero) | 17 |
| Putamen R | Subcortical | 16 |
| Postcentral Gyrus L | SomMot (Intero) | 15 |
| Cingulate Gyrus Post R | Cont (Intero) | 14 |
| Paracingulate Gyrus R | Cont (Intero) | 13 |
| Postcentral Gyrus R | SomMot (Intero) | 13 |
| Precentral Gyrus L | SomMot | 12 |
| Postcentral Gyrus R | SomMot (Intero) | 12 |
| Precentral Gyrus L | SomMot | 12 |
| Cingulate Gyrus Ant L | Default (Intero) | 11 |
| Precentral Gyrus R | SomMot | 11 |
| Precentral Gyrus R | SomMot | 11 |
| Caudate L | Subcortical | 10 |
| Superior Frontal Gyrus R | Cont | 9 |
| Precentral Gyrus L | SomMot | 8 |
| Precentral Gyrus L | SomMot | 8 |
| Precentral Gyrus R | SomMot | 7 |
| Postcentral Gyrus R | SomMot (Intero) | 7 |
| Cingulate Gyrus Post L | Cont (Intero) | 7 |
| Postcentral Gyrus L | SomMot (Intero) | 6 |
| Superior Parietal Lobule R | SomMot | 5 |
| Parietal Operculum Cortex R | SomMot | 5 |
| Postcentral Gyrus R | SomMot (Intero) | 4 |
| Central Opercular Cortex L | SomMot (Intero) | 4 |
| Insular Cortex R | SomMot (Intero) | 4 |
| Insular Cortex L | SomMot (Intero) | 4 |
| Juxtapositional Lobule Cortex R | SomMot | 4 |
| Juxtapositional Lobule Cortex L | SomMot | 4 |
| Middle Frontal Gyrus R | Cont | 4 |
| Precentral Gyrus L | SomMot | 3 |
| Middle Frontal Gyrus R | Cont | 3 |
| Frontal Pole R | Cont | 3 |
| Frontal Pole R | Cont | 2 |
| Planum Temporale R | SomMot | 2 |
| Superior Temporal Gyrus Ant R | Default | 2 |

| | | |
|---|---|---|
| Supramarginal Gyrus Post L | Default | 2 |
| Postcentral Gyrus L | DorsAttn (Intero) | 2 |
| Postcentral Gyrus L | DorsAttn (Intero) | 2 |
| Supramarginal Gyrus Post R | Cont | 2 |
| Lateral Occipital Cortex Inf L | Vis | 1 |
| Lingual Gyrus L | Vis | 1 |
| Superior Parietal Lobule L | SomMot | 1 |
| Central Opercular Cortex L | SomMot (Intero) | 1 |
| Planum Temporale L | SomMot | 1 |
| Cuneal Cortex L | Vis | 1 |
| Superior Frontal Gyrus L | SomMot | 1 |
| Central Opercular Cortex R | SomMot (Intero) | 1 |
| Temporal Occipital Fusiform Cortex L | DorsAttn | 1 |
| Superior Parietal Lobule L | DorsAttn | 1 |
| Superior Temporal Gyrus Post R | Default | 1 |
| Supramarginal Gyrus Post R | SalVentAttn | 1 |
| Superior Parietal Lobule R | DorsAttn | 1 |
| Lateral Occipital Cortex Sup R | DorsAttn | 1 |
| Postcentral Gyrus R | DorsAttn (Intero) | 1 |
| Postcentral Gyrus R | DorsAttn (Intero) | 1 |
| Middle Temporal Gyrus TempOcci R | DorsAttn | 1 |
| Inferior Temporal Gyrus TempOcci R | DorsAttn | 1 |
| Superior Frontal Gyrus R | SomMot | 1 |
| Precentral Gyrus R | SomMot | 1 |
| Lateral Occipital Cortex Inf R | Vis | 1 |
| Lingual Gyrus R | Vis | 1 |
| Temporal Fusiform Cortex Post R | Vis | 1 |
| Superior Temporal Gyrus Ant L | Default | 1 |
| Planum Temporale L | SalVentAttn | 1 |
| Lateral Occipital Cortex Inf L | Vis | 1 |

**Supplementary Table 8:** Significant connectivity edges from the permutation-based mass-univariate ordinary least squares analysis of the mind-wandering factors (see Figure 3 for the Exploratory Factor Analysis).

| Region 1 | Region 2 | Network 1 | Network 2 | t | *p* |
|---|---|---|---|---|---|
| | | **Factor 1** | | | |
| Lingual Gyrus L | Precuneous Cortex L | Vis | Cont | -5.348 | 0.002 |
| Occipital Fusiform Gyrus L | Frontal Pole L | Vis | Cont | -4.838 | 0.031 |
| Occipital Fusiform Gyrus L | Cingulate Gyrus Post R | Vis | Intero (SomMot) | -4.971 | 0.016 |
| Cuneal Cortex L | Middle Temporal Gyrus TempOcci R | Vis | SalVentAttn | 4.801 | 0.036 |
| Cuneal Cortex L | Precuneous Cortex R | Vis | Default | 4.939 | 0.019 |
| Cuneal Cortex L | Planum Polare L | Vis | SomMot | 4.798 | 0.037 |
| Superior Parietal Lobule L | Temporal Pole R | SomMot | Limbic | 5.546 | 0.001 |
| Superior Parietal Lobule L | Inferior Temporal Gyrus Post R | SomMot | Limbic | 5.836 | < .001 |
| Precentral Gyrus L | Cingulate Gyrus Post L | SomMot | Intero (Cont) | 5.874 | < .001 |
| Precentral Gyrus L | Cingulate Gyrus Ant L | SomMot | Intero (Cont) | 6.264 | < .001 |
| Temporal Occipital Fusiform Cortex L | Frontal Orbital Cortex L | DorsAttn | Intero (Default) | 4.881 | 0.026 |
| Lateral Occipital Cortex Sup L | Temporal Pole R | DorsAttn | Limbic | 4.732 | 0.050 |
| Middle Frontal Gyrus L | Precuneous Cortex R | Default | Cont | 5.155 | 0.006 |
| Central Opercular Cortex R | Superior Frontal Gyrus R | Intero (SalVentAttn) | Cont | 4.773 | 0.041 |
| Central Opercular Cortex R | Lateral Occipital Cortex Sup R | Intero | Default | 4.877 | 0.026 |

| | | | | | |
|---|---|---|---|---|---|
| | | (SalVentAttn) | | | |
| Cingulate Gyrus Ant R | Pallidum L | Intero (SalVentAttn) | Subcortical | 4.838 | 0.031 |
| Cingulate Gyrus Ant R | Putamen R | Intero (SalVentAttn) | Subcortical | 4.761 | 0.043 |
| | | **Factor 2** | | | |
| Lateral Occipital Cortex Sup R | Pallidum R | Default | Subcortical | 4.732 | 0.047 |

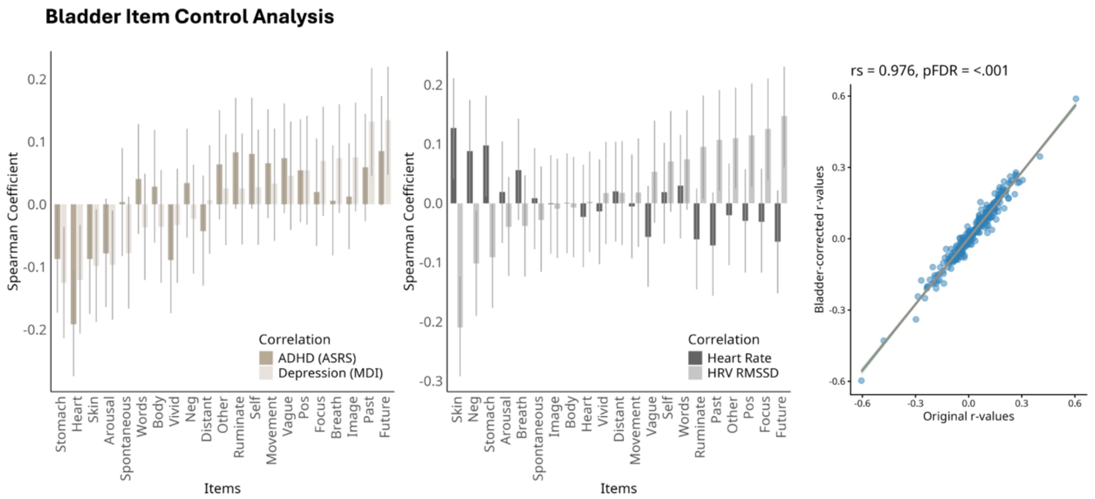


***Supplementary Figure 7. Robustness of psychophysiological associations after controlling for bladder-related thoughts.***

To determine if results were driven by specific interoceptive content, individual ratings for the item "My thoughts were about my bladder" were regressed out of all other mind-wandering items prior to correlational analysis. (Left and Middle) Spearman correlation coefficients between bladder-corrected mind-wandering items and psychophysiological variables. The overall pattern of associations remains highly stable. Core relationships (ADHD-Heart, Depression-Heart/Stomach/Future/Past, HeartRate-Skin, and HRV-Skin/Future/Past/Other/Focus/Pos) remain significant, while a subset of weaker associations (ADHD-Skin, Depression-Arousal/Skin, HeartRate-Neg, HRV-Neg) are no longer significant, indicating they were partially influenced by shared variance with bladder-related thoughts. (Right) Scatterplot comparing the original r-values (x-axis) with the bladder-corrected r-values (y-axis). The high correlation ($r_s = 0.976$, $p_{FDR} < .001$) confirms that the reported psychophysiological fingerprints are not artifacts of bladder attention.

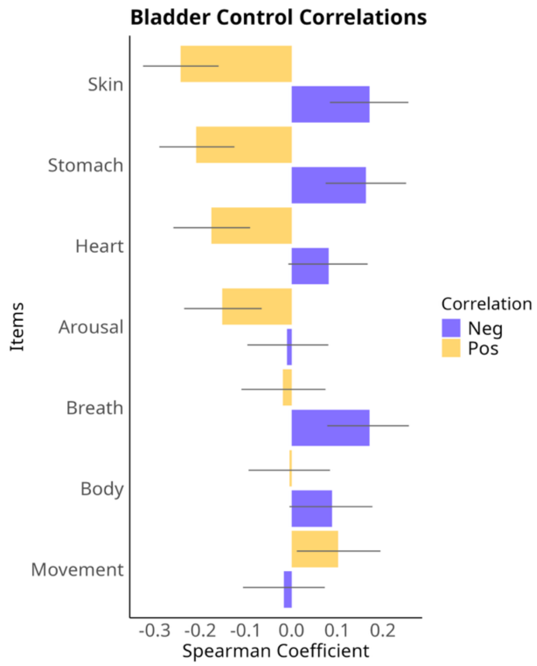


***Supplementary Figure 8. Robustness of affect-body associations after controlling for bladder awareness.***

Individual ratings for the item "My thoughts were about my bladder" were regressed from the mind-wandering items before correlating them with the items "My thoughts were positive" and "My thoughts were negative." Spearman correlation coefficients after correction (Purple = Negative; Yellow = Positive). The overall pattern of association between body items and affect was shared between the original and control analyses: somatic thoughts generally correlates with reduced ratings of "Positive" thoughts (Yellow: Skin, Stomach, Heart, Arousal) and increased ratings of "Negative" thoughts (Purple: Skin, Stomach, Breath). However, specific correlations between "Negative" thoughts and Heart/Body, as well as "Positive" thoughts and Movement, did not reach significance when controlling for bladder oriented thoughts.

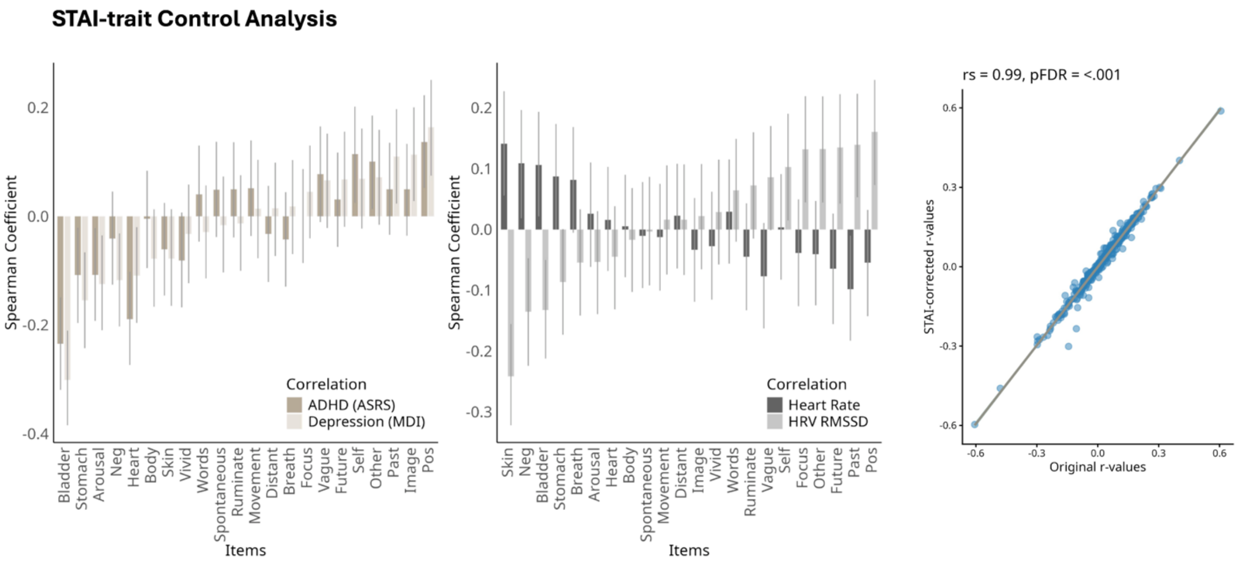


***Supplementary Figure 9. Robustness of psychophysiological associations after controlling for trait anxiety (STAI).*** To determine if the link between body-wandering and mental health/physiology was driven by anxiety predisposition, individual Trait Anxiety (STAI-T) scores were regressed out of all mind-wandering items prior to correlational analysis. (Left and Middle) Spearman correlation coefficients between anxiety-corrected mind-wandering items and psychophysiological variables. The overall pattern of associations remains highly stable. Key relationships (ADHD-Heart/Bladder, Depression-Arousal/Heart/Bladder/Stomach/Past, HeartRate-Bladder/Skin/Neg, and HRV-Bladder/Skin/Future/Past/Other/Focus/Pos/Neg) remain significant. While a small subset of peripheral associations (ADHD-Skin, Depression-Skin/Future) did not survive correction, new associations emerged (ADHD-Arousal/Stomach/Self/Pos, Depression-Image/Pos/Neg, HRV-Self), suggesting that removing unique anxiety-related variance may actually unmask specific somatosensory connections. (Right) Scatterplot comparing the original r-values with the STAI-corrected r-values. The correlation pattern largely persists, supporting that the "body-wandering" phenotype is distinct from trait anxiety.

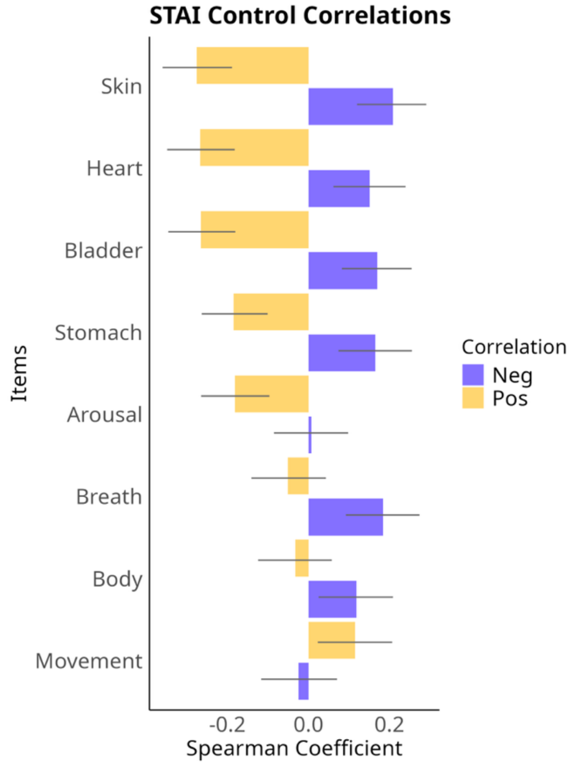


***Supplementary Figure 10*****: Spearman correlation analysis of affect and body-related mind-wandering items once controlled for individual differences in trait anxiety (STAI) ratings.**

To determine if the link between body-focused thought and negative affect was driven by trait anxiety, individual STAI-Trait scores were regressed from all mind-wandering items. The residuals were correlated with "My thoughts were positive" and "My thoughts were negative." All significant spearman correlation coefficients persisted after correction. The analysis reveals a pattern highly similar to the original uncorrected analysis: body-related items (Heart, Skin, Stomach, Breath, Body) remain significantly associated with higher "Negative" ratings and/or lower "Positive" ratings. This persistence confirms that the negative affective tone of body-wandering is a core feature of the embodied thought state, rather than an artifact of high trait anxiety.

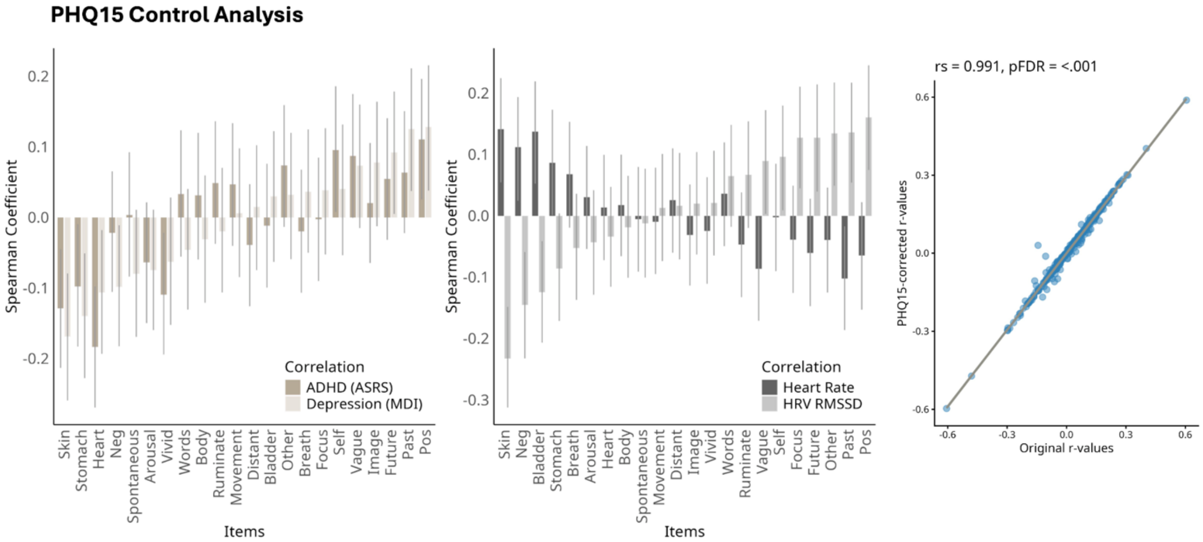


***Supplementary Figure 11. Robustness of psychophysiological associations after controlling for trait somatic symptoms (PHQ-15).***

To account for potential confounding by clinical somatic symptoms, total PHQ-15 scores were regressed out of mind-wandering items prior to correlational analysis. (Left and Middle) Spearman correlations between PHQ-corrected mind-wandering items and psychophysiological variables. The overall pattern of associations remains highly stable. Key relationships remain significant after correction, including: ADHD-Heart/Skin; Depression-Heart/Stomach/Skin/Past; Heart Rate-Bladder/Skin/Neg; and HRV-Bladder/Skin/Future/Past/Other/Focus/Pos/Neg. A subset of associations (ADHD-Bladder, Depression-Arousal/Bladder/Future) did not reach significance after correction, suggesting these specific effects were influenced by trait-level somatic symptoms. (Right) Scatterplot comparing the original r-values (x-axis) with the PHQ-corrected r-values (y-axis). The very high correlation (rs = 0.991, pFDR < .001) confirms that the reported psychophysiological fingerprints are robust from trait-level somatic symptom severity.

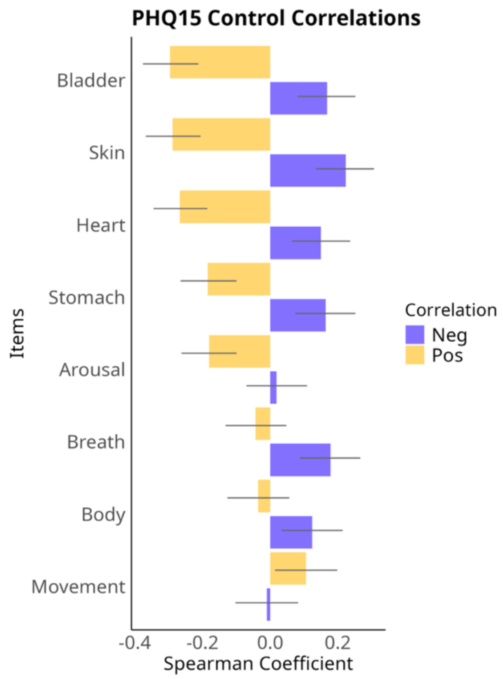


***Supplementary Figure 12. Robustness of affect-body associations after controlling for trait somatic symptoms (PHQ-15).***

To determine if the relationship between affective valence and body-focused thought was driven by clinical somatic symptoms, total PHQ-15 scores were regressed from all mind-wandering items. The resulting residuals were correlated with the items "My thoughts were positive" and "My thoughts were negative." Spearman correlation coefficients after correction (Purple = Negative; Yellow = Positive). The results reveal a pattern highly similar to the original uncorrected analysis in Figure 1, with all significant correlations persisting from before and after PHQ15-correction. Specifically, body-related items (Bladder, Skin, Heart, Stomach, Breath, Body) remain significantly associated with higher "Negative" ratings and/or lower "Positive" ratings. The persistence of these correlations indicates that the affective signature of body-wandering is not explained by trait-level somatic symptom severity.

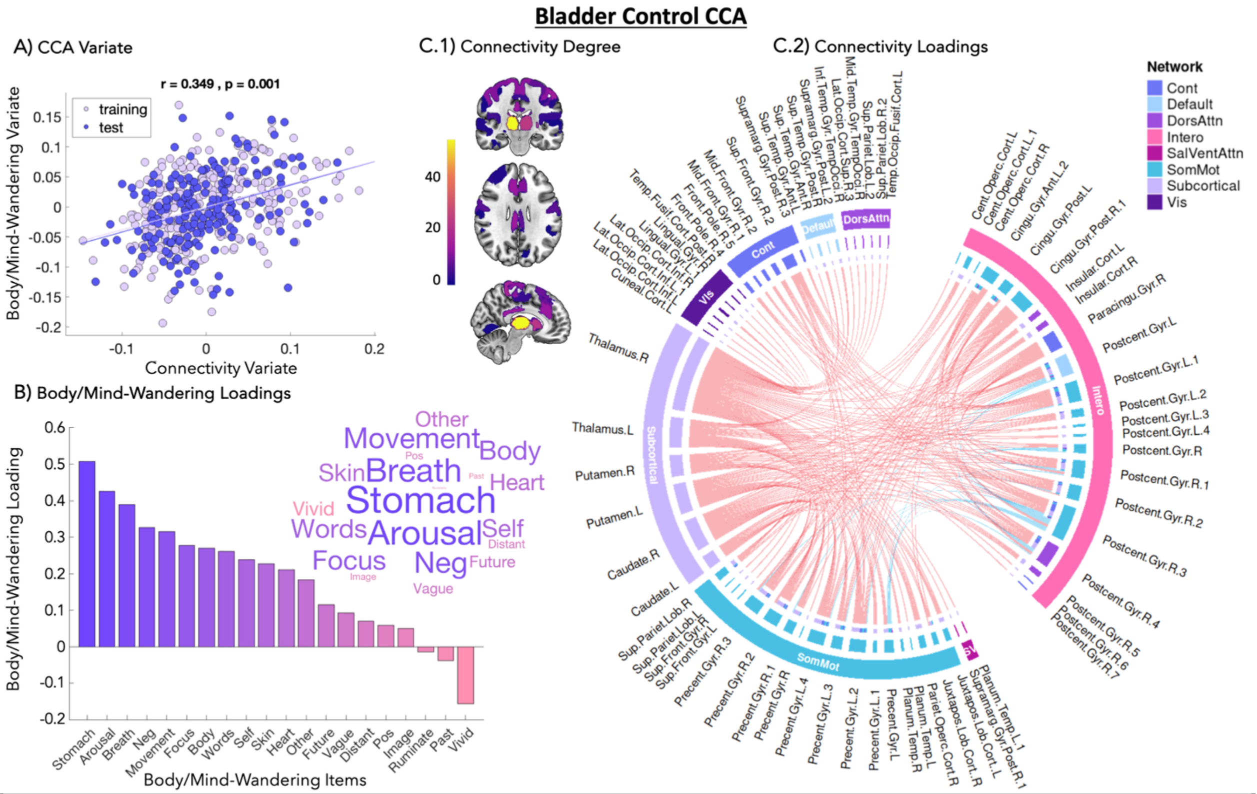


***Supplementary Figure 13*****: Body/mind-wandering and brain connectivity CCA with the 'Bladder' item included as a nuisance regressor.**

To determine if the multivariate relationship between mind-wandering and functional connectivity was driven by visceral discomfort, the Canonical Correlation Analysis (CCA) was re-computed with the item "My thoughts were about my bladder" included as a nuisance regressor. (A) The canonical correlation remains statistically significant ($r = 0.349$, $p = 0.001$). (B) Thought item loadings. The composition of the thought variate is highly similar to the original analysis, characterized by positive loadings for body-focused items (Stomach, Arousal, Breath) and Negative affect. (C) Connectivity loadings. The neural signature remains stable, dominated by connectivity within and between Interoceptive, Somatomotor, and Subcortical networks. These results support that the identified cortical-subcortical signature of embodied mind-wandering is not an artifact of bladder attention.

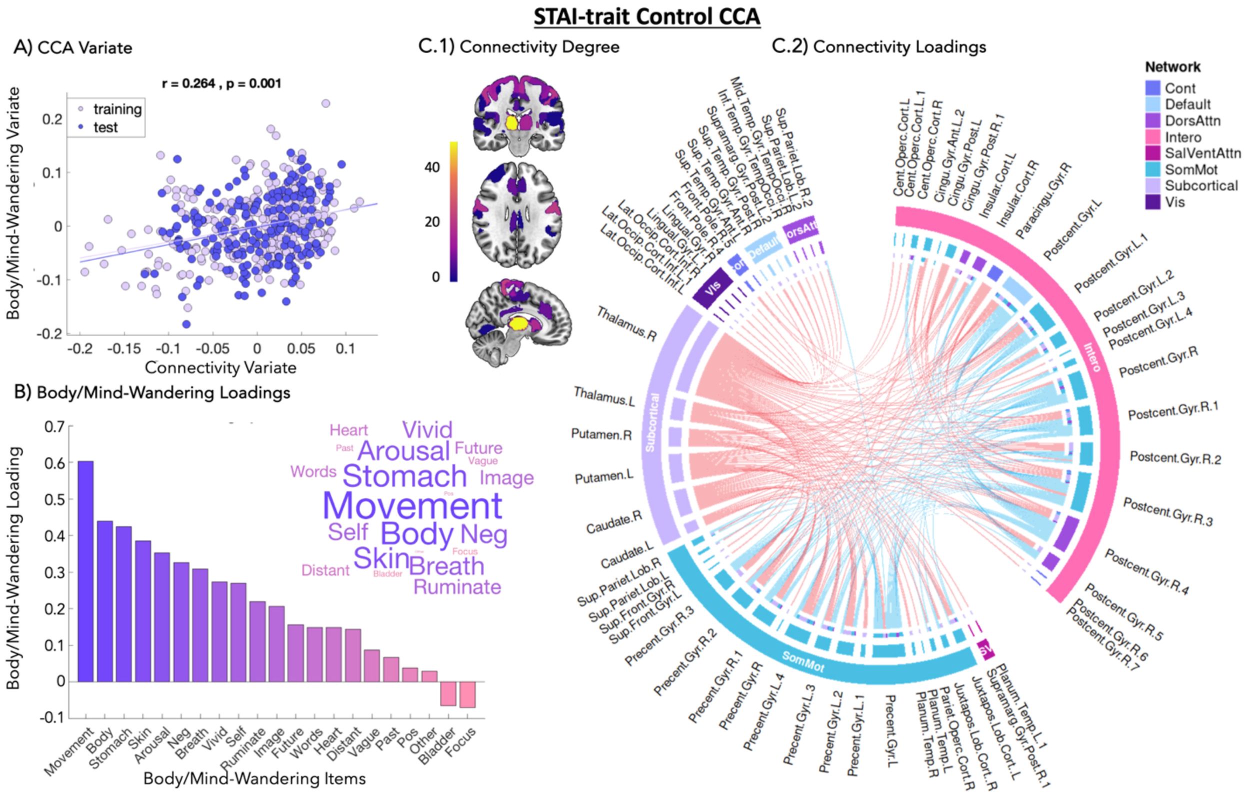


#### *Supplementary Figure 14. Robustness of the brain-body neural signature after controlling for Trait Anxiety.*

To ensure the multivariate relationship between mind-wandering and functional connectivity was not driven by anxious arousal, the Canonical Correlation Analysis (CCA) was re-calculated with STAI-Trait scores included as a nuisance regressor. (A) The canonical correlation remains statistically significant ($r = 0.264$, $p = 0.001$). (B & C) Both the thought loadings (B) and connectivity patterns (C) are highly consistent with the original analysis. The persistence of the Interoceptive, Somatomotor, and Subcortical connectivity profile suggests that individual differences in trait anxiety are not a major factor in the neural signature of embodied mind-wandering.

**PHQ15 Control CCA**

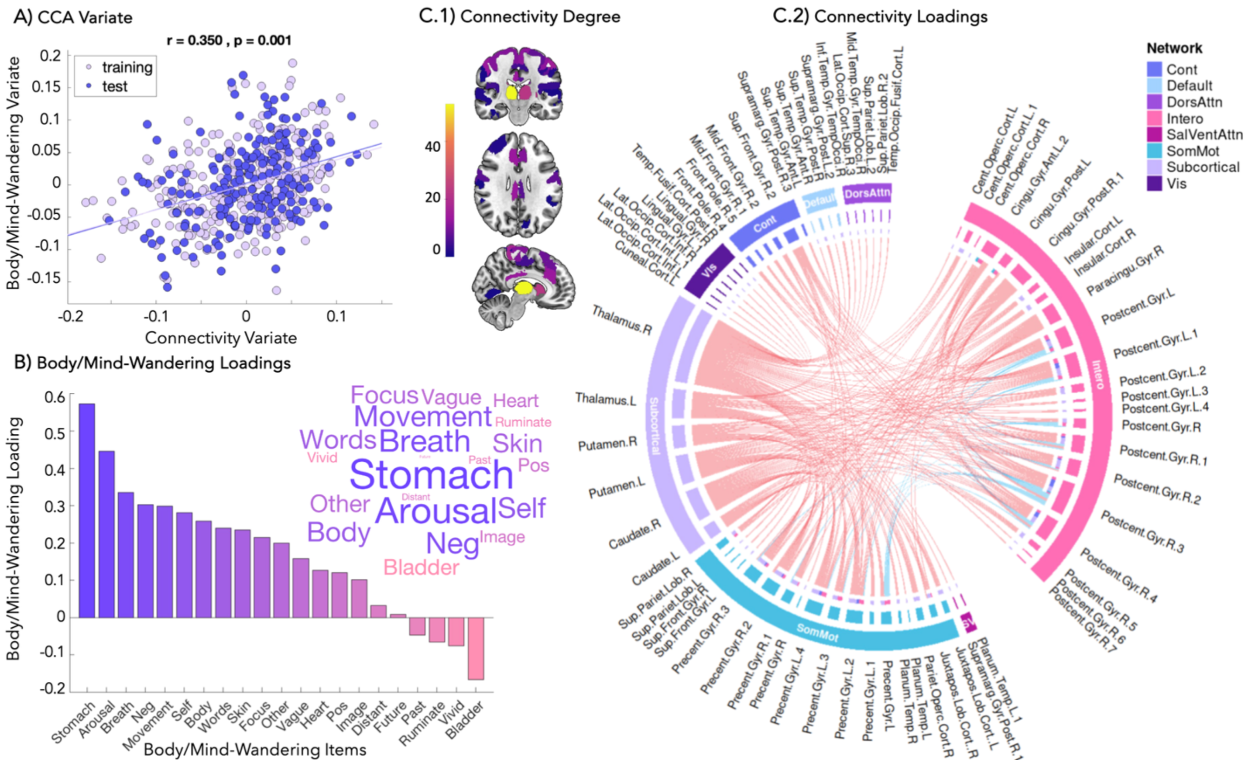


***Supplementary Figure 15. Robustness of the brain-body neural signature after controlling for trait somatic symptoms.***

To determine if the multivariate relationship between mind-wandering and functional connectivity was driven by clinical somatic symptoms, the Canonical Correlation Analysis (CCA) was re-computed with PHQ-15 sum-scores (Kroenke et al., 2002) included as a nuisance regressor. (A) The canonical correlation remains statistically significant ($r = 0.350$, $p = 0.001$). (B & C) Both the thought loadings and connectivity patterns are highly consistent with the original analysis presented in the main text. The persistence of the Interoceptive, Somatomotor, and Subcortical connectivity profile supports that the neural signature of embodied mind-wandering is not driven by trait-level somatic symptom severity.

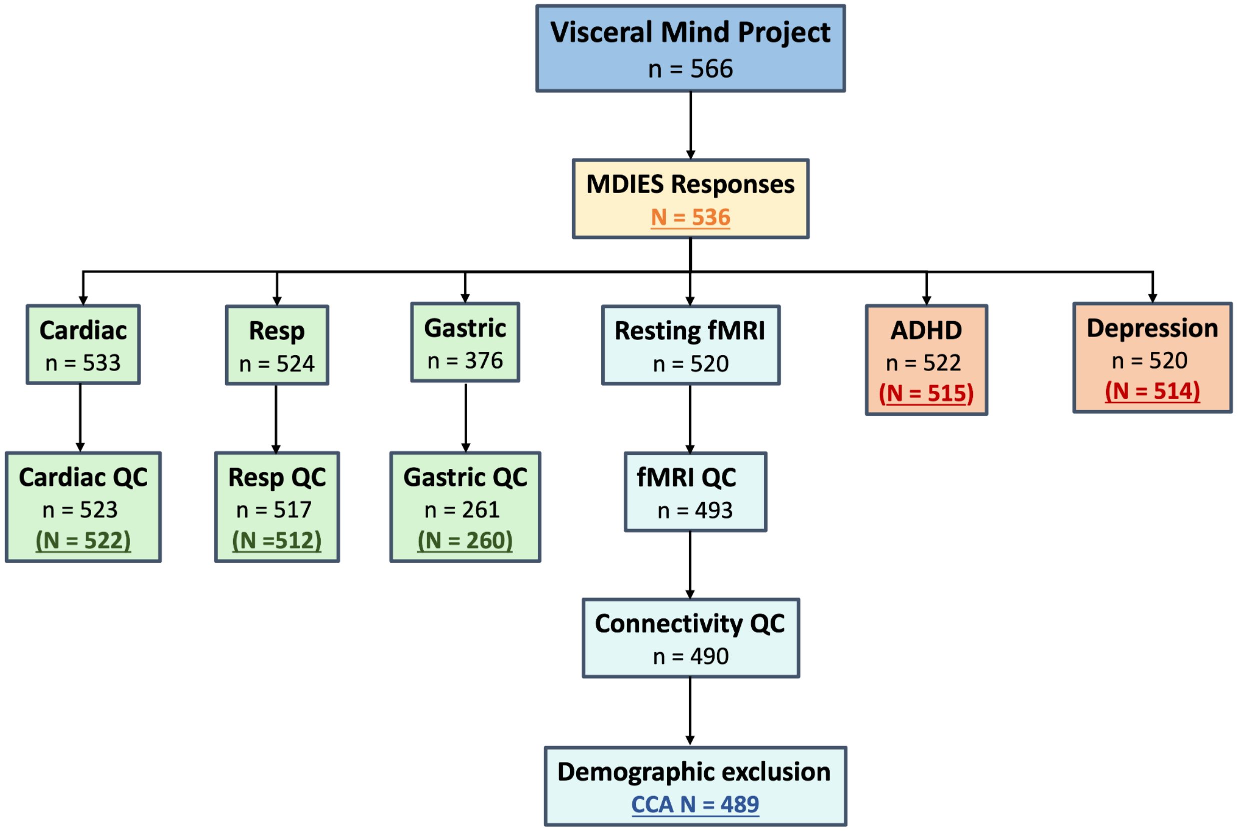


***Supplementary Figure 16*: Flow chart of analysis sample sizes from the Visceral Mind Project.**

Flow chart describing the sample size of each data type from the Visceral Mind Project, including those who completed the post-resting-fMRI Multi-Dimensional Interoceptive Experience Sampling (MDIES) of mind-wandering experiences (in yellow: N=536). Other data types used for further analyses include resting-state fMRI (in blue), physiological recordings (in green: including cardiac photoplethysmography, respiratory breathing belt, and gastric electrogastrography), and ADHD and Depression scores (in orange). The neuroimaging and physiological data were visually inspected resulting in low quality check (QC) data rejections. The sample for estimating the functional connectivity fingerprints of embodied mind-wandering via canonical correlation analysis (CCA) was 489. Outliers were rejected from the physiological data and the survey scores (in brackets: outliers were computed as 3 standard deviations from the mean). The final sample for the physiological and psychological fingerprints of body-wandering and cognitive-wandering are shown underlined in bold respectively.

### Supplementary fMRI preprocessing workflow

#### fMRIprep preprocessing

We implemented the minimal preprocessing pipeline in fmriprep. fMRI results included in this manuscript come from preprocessing performed using fMRIPrep 22.1.1 ((Esteban et al., 2019, 2022); RRID:SCR_016216), which is based on Nipype 1.8.5 ((Gorgolewski et al., 2018); RRID:SCR_002502).

#### Anatomical data preprocessing

A total of 1 T1-weighted (T1w) images were found within the input BIDS dataset. The T1-weighted (T1w) image was corrected for intensity non-uniformity (INU) with N4BiasFieldCorrection (Tustison et al., 2010), distributed with ANTs 2.3.3 ((Avants et al., 2008), RRID:SCR_004757), and used as T1w-reference throughout the workflow. The T1w-reference was then skull-stripped with a Nipype implementation of the antsBrainExtraction.sh workflow (from ANTs), using OASIS30ANTs as target template. Brain tissue segmentation of cerebrospinal fluid (CSF), white-matter (WM) and gray-matter (GM) was performed on the brain-extracted T1w using FAST (FSL 6.0.5.1:57b01774, RRID:SCR_002823, (Zhang et al., 2001)). Brain surfaces were reconstructed using recon-all (FreeSurfer 7.2.0, RRID:SCR_001847, (Dale et al., 1999)), and the brain mask estimated previously was refined with a custom variation of the method to reconcile ANTs-derived and FreeSurfer-derived segmentations of the cortical gray-matter of Mindboggle (RRID:SCR_002438, (Klein et al., 2017)). Volume-based spatial normalization to two standard spaces (MNI152NLin2009cAsym, MNI152NLin6Asym) was performed through nonlinear registration with antsRegistration (ANTs 2.3.3), using brain-extracted versions of both T1w reference and the T1w template. The following templates were selected for spatial normalization: ICBM 152 Nonlinear Asymmetrical template version 2009c [(Fonov et al., 2009), RRID:SCR_008796; TemplateFlow ID: MNI152NLin2009cAsym], FSL's MNI ICBM 152 non-linear 6th Generation Asymmetric Average Brain Stereotaxic Registration Model [(Evans et al., 2012), RRID:SCR_002823; TemplateFlow ID: MNI152NLin6Asym].

#### Functional data preprocessing

For each of the 3 BOLD runs found per subject (across all tasks and sessions), the following preprocessing was performed. First, a reference volume and its skull-stripped version were generated using a custom methodology of fMRIPrep. Head-motion parameters with respect to the BOLD reference (transformation matrices, and six corresponding rotation and translation parameters) are estimated before any spatiotemporal filtering using mcflirt(FSL 6.0.5.1:57b01774, (Jenkinson et al., 2002)). BOLD runs were slice-time corrected to 0.658s (0.5 of slice acquisition range 0s-1.31s) using 3dTshift from AFNI ((Cox & Hyde, 1997), RRID:SCR_005927). The BOLD time-series (including slice-timing correction when applied) were resampled onto their original, native space by applying the transforms to correct for head-motion. These resampled BOLD time-series will be referred to as preprocessed BOLD in original space, or just preprocessed

BOLD. The BOLD reference was then co-registered to the T1w reference using bbregister (FreeSurfer) which implements boundary-based registration (Greve & Fischl, 2009). Co-registration was configured with six degrees of freedom. Several confounding time-series were calculated based on the preprocessed BOLD: framewise displacement (FD), DVARS and three region-wise global signals. FD was computed using two formulations following Power (absolute sum of relative motions, (Power et al., 2014)) and Jenkinson (relative root mean square displacement between affines, (Jenkinson et al., 2002)). FD and DVARS are calculated for each functional run, both using their implementations in Nipype (following the definitions by (Power et al., 2014)). The three global signals are extracted within the CSF, the WM, and the whole-brain masks. Additionally, a set of physiological regressors were extracted to allow for component-based noise correction (CompCor, (Behzadi et al., 2007)). Principal components are estimated after high-pass filtering the preprocessed BOLD time-series (using a discrete cosine filter with 128s cut-off) for the two CompCor variants: temporal (tCompCor) and anatomical (aCompCor). tCompCor components are then calculated from the top 2% variable voxels within the brain mask. For aCompCor, three probabilistic masks (CSF, WM and combined CSF+WM) are generated in anatomical space. The implementation differs from that of Behzadi et al. in that instead of eroding the masks by 2 pixels on BOLD space, a mask of pixels that likely contain a volume fraction of GM is subtracted from the aCompCor masks. This mask is obtained by dilating a GM mask extracted from the FreeSurfer's aseg segmentation, and it ensures components are not extracted from voxels containing a minimal fraction of GM. Finally, these masks are resampled into BOLD space and binarized by thresholding at 0.99 (as in the original implementation). Components are also calculated separately within the WM and CSF masks. For each CompCor decomposition, the k components with the largest singular values are retained, such that the retained components' time series are sufficient to explain 50 percent of variance across the nuisance mask (CSF, WM, combined, or temporal). The remaining components are dropped from consideration. The head-motion estimates calculated in the correction step were also placed within the corresponding confounds file. The confound time series derived from head motion estimates and global signals were expanded with the inclusion of temporal derivatives and quadratic terms for each (Satterthwaite et al., 2013). Frames that exceeded a threshold of 0.5 mm FD or 1.5 standardized DVARS were annotated as motion outliers. Additional nuisance timeseries are calculated by means of principal components analysis of the signal found within a thin band (crown) of voxels around the edge of the brain, as proposed by (Patriat et al., 2017). The BOLD time-series were resampled into standard space, generating a preprocessed BOLD run in MNI152NLin2009cAsym space. First, a reference volume and its skull-stripped version were generated using a custom methodology of fMRIPrep. The BOLD time-series were resampled onto the following surfaces (FreeSurfer reconstruction nomenclature): fsaverage. Grayordinates files (Glasser et al., 2013) containing 91k samples were also generated using the highest-resolution fsaverage as intermediate standardized surface space. All resamplings can be performed with a single interpolation step by composing all the pertinent transformations (head-motion transform matrices, susceptibility distortion correction when available, and co-registrations to anatomical and output spaces).

Gridded (volumetric) resamplings were performed using antsApplyTransforms (ANTs), configured with Lanczos interpolation to minimize the smoothing effects of other kernels (Lanczos, 1964). Non-gridded (surface) resamplings were performed using mri_vol2surf (FreeSurfer).

Many internal operations of fMRIPrep use Nilearn 0.9.1 ((Abraham et al., 2014), RRID:SCR_001362), mostly within the functional processing workflow. For more details of the pipeline, see (*Preprocessing Pipeline Details - Fmriprep Version Documentation*).

#### Functional connectivity

Preprocessing following fMRIPrep included extracting a parcellated time-series using a 200-cortical-region Schaefer atlas with an additional 16 subcortical regions (Schaefer et al., 2018). This was performed using the Nilearn functions "NiftiMapsMasker" and "fit_transform" which additionally regressed out noise parameters estimated from fMRIPrep (24 motion parameters, two white matter and cerebrospinal fluid noise parameters, a high pass filter, and six aCompCor 'anat_combined' parameters (Abraham et al., 2014; Behzadi et al., 2007)). We also low pass filtered at 0.08. Finally, we estimated functional connectivity for each participant using a full 216-node correlation matrix via the Nilearn "ConnectivityMeasure(kind=correlation).fit_transform" function.

### Supplementary physiological data preprocessing workflow

#### Physiological recording acquisition

We simultaneously recorded physiological measurements (photoplethysmography, respiratory breathing belt, and electrogastrography) during resting-state fMRI. The photoplethysmography was recorded on the right cheek via a Sentec Digital Monitoring system for the first data collection cohort at 1000 Hz (208 participants) and on the left index/middle finger via the MRI scanner for the second cohort at 200 Hz (325 participants). We obtained respiratory recordings using a Siemens respiratory cushion and belt system. The first data collection cohort was recorded via a Brain Vision ExG system at 1000 Hz (208 participants), while the second cohort was recorded via an MRI scanner at 50 Hz (316 participants). For the EGG recordings, we cleaned the abdomen and applied abrasive gel to remove dead skin and improve the signal-to-noise ratio. Three electrogastrography recording montages (1, 3 or 6 bipolar channels) were implemented using a Brain Vision MRI-compatible ExG system and amplifier. All ExG physiological recordings were acquired with a sampling rate of 1000 Hz, a low-pass filter of 1000 Hz (with a 450 Hz anti-aliasing filter), and no high-pass filter (DC recordings). EGG was recorded at a 0.5 µV/bit resolution, and +/- 16.384 mV range, while photoplethysmography and respiratory recordings were acquired at 152.6 µV/bit resolution, and +/- 5000 mV range.

#### Cardiac preprocessing

R-peaks were identified from the cardiac recordings using the python package ‘systole’ using the ‘rolling_average_ppg’ method, while correcting for clipping artefacts in the second cohort’s data using a threshold of 950’000 and a cubic spline interpolation method. All the detected R-peaks were visually inspected and manually corrected, if necessary, as well as unintelligible segments marked and discarded. We excluded participants with less than 2 minutes of continuous cardiac recording. We computed heart rate as the mean of the beats per minute, while heart rate variability was defined as the root mean square of successive differences of the R-R intervals in milliseconds (RMSSD). Heart rate variability in the frequency domain was calculated by estimating a Welch power spectrum density. The high frequency power in normalised units was defined as high frequency power (0.15 Hz - 0.4 Hz), divided by high and low frequency combined (0.04 Hz - 0.4 Hz), multiplied by 100 (Legrand & Allen, 2022).

#### Respiratory preprocessing

To address occasional missing samples from the scanner respiratory recordings, we performed linear interpolation on the second cohort's data. We excluded respiratory data if the recording was less than 7 minutes. We preprocessed the respiratory recordings by Butterworth filtering with a cutoff of 1 and an order of 4. We computed the z-score of the respiratory recordings for peak (i.e., inhalation) and trough (i.e., exhalation) detection. For inhalation peak detection, we employed the 'find_peaks' function from the ‘scipy.signal’ Python library, setting distance to 1, prominence to

0.6, and width to 2. We defined exhalation troughs as the minimum points between the identified peaks. We visually inspected peak and trough detection, manually correcting, if necessary, by adding missing peaks/troughs or removing incorrect ones.

Breath duration was calculated as the time between peaks in seconds, inspiration depth as the difference in signal strength between the peak and the preceding trough (i.e., peak minus previous trough), and expiration depth as the difference in signal strength between the peak and the subsequent trough (i.e., peak minus following trough). We computed the mean and standard deviation of these single peak-to-peak and trough measurements, yielding the following respiratory metrics: mean breath duration, breath duration standard deviation, mean breath depth, and breath depth standard deviation (averaging/standard deviation across inspiration and expiration depths). Additionally, we defined the respiration rate as the number of breaths per minute.

#### Gastric preprocessing

The EGG data was first demeaned and downsampled from 1000 Hz to 10 Hz for computational efficiency, followed by computing the power spectrum using a Hanning-tapered fast Fourier transform incorporating 1000 seconds of zero-padding in 200-second data segments with 75% overlap. For each participant, we selected the bipolar EGG channel that showed the most prominent peak within the normogastric range (0.033-0.066 Hz). Specifically, two independent researchers conducted peak selection by visually inspecting each channel to identify the EGG channel with the highest normogastric power peak, without large artefacts and with power above 5 µv2. Peak quality was rated as 'excellent' for Gaussian-like peaks (n=184) and 'good' for shoulder-like peaks (n=81); those not meeting these standards were deemed 'poor quality' (n=115) and excluded. This visual inspection approach is consistent with previous research to account for noise in the normogastric window, or cases of multiple peaks (Müller et al., 2022; Rebollo et al., 2018; Rebollo & Tallon-Baudry, 2021). As an additional check, we computed signal quality metrics using a comparison template-based procedure of 10 ideal participants with very clear and prominent gastric peaks. 'Poor quality' participants had significantly lower signal quality as measured by cosine similarity (excellent/good quality: Median = 0.963, Range = 0.667, poor quality: Median = 0.595, Range = 0.585; U = 63780, $p < .001$, rrb = 3.186) and Pearson's correlation (excellent/good quality: Median = 0.950, Range = 1.169, poor quality: Median = 0.054, Range = 1.287; U = 63849, $p < 0.001$, rrb = 3.190) (see Supplementary Figure 17). The selected EGG channel was then bandpass filtered, centred at the individual peak frequency (filter width of ±0.015 Hz, filter order of 5 or 1470 samples), in forward and backward direction to avoid time shifts.

From the computed EGG power spectra, we quantified the following normogastric EGG metrics: peak frequency, maximum power and proportion of power. Specifically, within the normogastric frequency range (0.033-0.067 Hz/2-4 cpm/15-30 seconds), we stored the peak frequency and maximum power. Furthermore, we computed the proportion of normogastric power as the sum of the normogastric power divided by the sum of the power in all gastric frequencies (including bradygastric, normogastric and tachygastric frequencies: 0.02-0.17 Hz/1-10 cpm/6-60 seconds).

#### Physiological metric processing

We log transformed the following physiological metrics to ensure approximate normal distributions: RMSSD, LF/HF ratio, breathing duration variability, maximum power in normogastria. Furthermore, we removed outliers from all physiological metrics as 3 standard deviations from the mean.

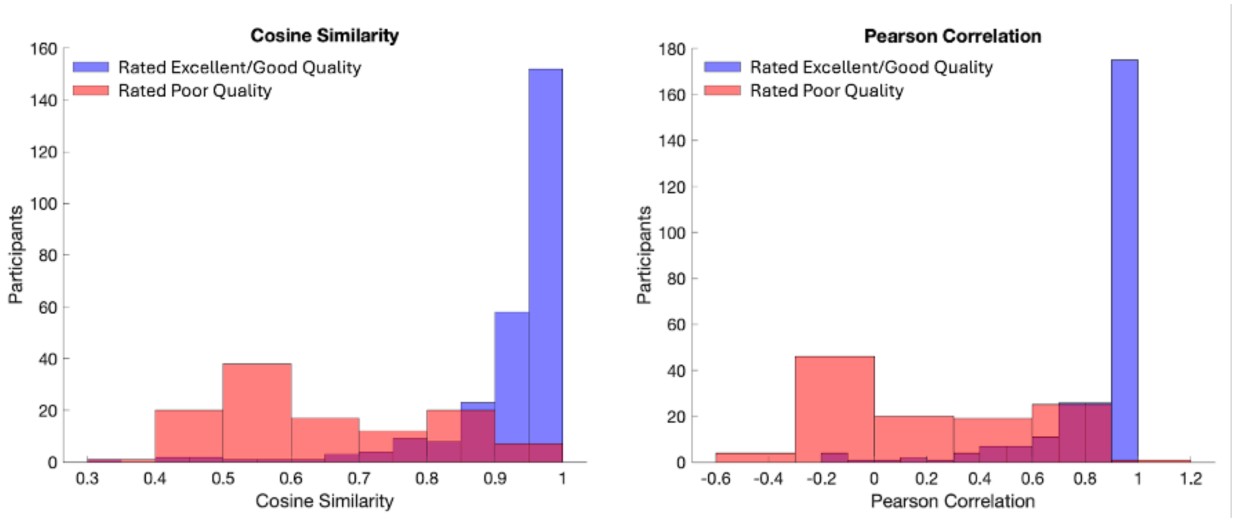


***Supplementary Figure 17:*** **Electrogastrography signal quality of included and excluded participants.**

EGG signal quality metrics (left: cosine similarity, right: Pearson correlation) of participants rated as ‘excellent/good quality’ in red and those rated as ‘poor quality’ in purple. These signal quality metrics were computed by comparing each individual's gastric FFT with the average FFT of 10 ideal participants with a very clear prominent normogastric peak.

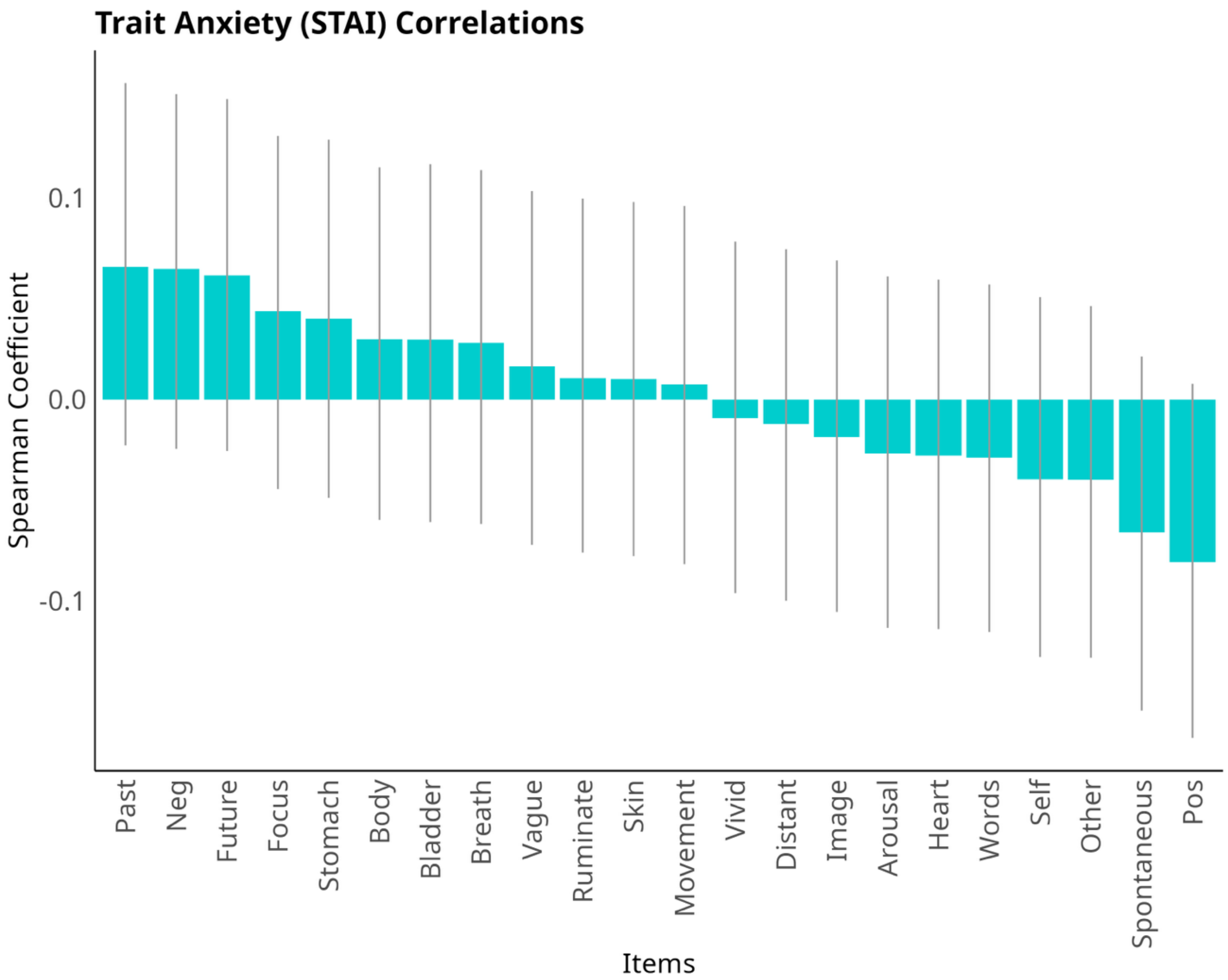


***Supplementary Figure 18:*** **Mind-wandering item correlations with trait anxiety sum scores (STAI-trait survey).** Spearman correlation coefficients of trait anxiety (STAI-trait survey sum score) with mind-wandering items, revealing no significant relationships. Error bars reflect bootstrap 95% confidence intervals via 10’000 resamples.
